# Extrinsic cues unlock cross-germ layer differentiation potential of CNS stem cells during regeneration

**DOI:** 10.1101/2025.11.24.690179

**Authors:** Civia Z. Chen, Yizhou Yu, Natalia Murphy, Juan F. Cubillos, Khalil S. Rawji, Chao Zhao, Myfanwy Hill, Peter Arthur-Farraj, Robin J.M. Franklin, Björn Neumann

## Abstract

During CNS regeneration, neuroepithelial-derived oligodendrocyte progenitor cells (OPCs) can cross germ-layer boundaries to generate neural-crest-derived Schwann cells (SCs). However, the underlying mechanism and disease relevance of this unique phenomenon of cellular plasticity is unclear. Here, we combine single cell genomics in rodent models of CNS injury and samples from multiple sclerosis patients to characterise OPC-derived SCs. We discover that integrin signalling and bone morphogenetic protein (BMP) activation activate a core SOX10/OLIG2 transcriptional circuit to drive OPC-to-SC differentiation. We show that OPC-derived SCs myelinate *in vitro* and *in vivo*, and unlike peripheral SCs, can integrate into astrocyte-rich territories. Together, these findings define a conserved molecular mechanism of adult plasticity, enabling the control of OPC fate choices beyond their germ layer origin for CNS repair.

## INTRODUCTION

The anatomical segregation of the vertebrate nervous system into central (CNS) and peripheral (PNS) compartments is among the most rigid boundaries established during development. This division is evident at the nerve root transition zones, where myelination shifts from neuroepithelial-derived oligodendrocytes to neural-crest-derived Schwann cells (SCs). The boundary between the CNS and PNS is actively maintained by astrocytes which physically acts as a non-permissive barrier and secretes inhibitory cues to exclude peripheral SCs from the spinal cord or brain^1^. Although SC transplantation remains one of the most promising strategies for promoting CNS axonal regeneration, astrocytic inhibition is the primary obstruction for wider use of SC transplantation in CNS regenerative medicine^2,3^.

In vertebrate species, differentiation is typically restricted to the cell types of their parent lineage. However, the adult CNS harbours an extreme capacity for lineage plasticity. Following demyelination or traumatic injury, OPCs become activated, migrate, proliferate and differentiate into a new myelinating cell. In most cases, OPCs give rise to oligodendrocytes, as expected. However, in certain situations, notably when there is concurrent loss of astrocytes, a subset of OPCs will generate Schwann cells (SCs)^4^, the myelinating cells of the peripheral nervous system and a cell of neural crest origin. CNS remyelination by SCs is seen in clinical disease, such as multiple sclerosis (MS)^5,6^ and traumatic CNS injury^7,8^, as well as in a range of experimental demyelination models^9–12^. The neural crest is now regarded as the “fourth” germ layer^13^, making the differentiation of the neuroepithelium-derived OPC into a cell of neural crest origin a unique cross-germ layer event during tissue regeneration. However, the mechanisms enabling this cross-germ layer transition are unknown.

Existing models suggest OPC-to-SC transitions are driven by high levels of BMP signalling that is unopposed by the presence or activity of astrocytes in the lesion environment^14,15^. However, BMP signalling alone is insufficient to induce this fate transition in pure cultures of isolated OPCs *in vitro*. Here, we identify the injury-associated extracellular matrix (ECM) glycoprotein vitronectin as a critical signal that stabilises a SOX10-dependent program to override the default OLIG2-driven oligodendrocyte identity. We demonstrate that this mechanism is a conserved feature in human disease and show that direct manipulation of the SOX10/OLIG2 transcriptional checkpoint is sufficient to overcome astrocytic-mediated inhibition to allow OPCs to differentiate into SCs (which we term oSCs). Our results identify the SOX10/OLIG2 circuit as the fundamental gatekeeper of neuroepithelial-to-neural crest plasticity and establish a strategy to generate CNS SCs that can successfully integrate into the otherwise exclusionary, astrocyte-rich CNS environment for repair.

## RESULTS

### Transcriptomic profiling of OPCs activated by demyelination reveals a Schwann cell-primed subpopulation

To characterise the transcriptomic landscape of OPCs at the onset of differentiation, we performed Smartseq2 single cell transcriptomics on focal ethidium-bromide-induced demyelinating lesions in the rat caudal cerebellar peduncle (CCP) at 10 days post-injection – a key temporal window where OPCs recruited into the lesion are beginning to undergo differentiation^16^. This yielded 517 high-quality cells across three biological replicates (Fig. 1a, Extended Data Fig. 1a, b).

**Fig. 1:**
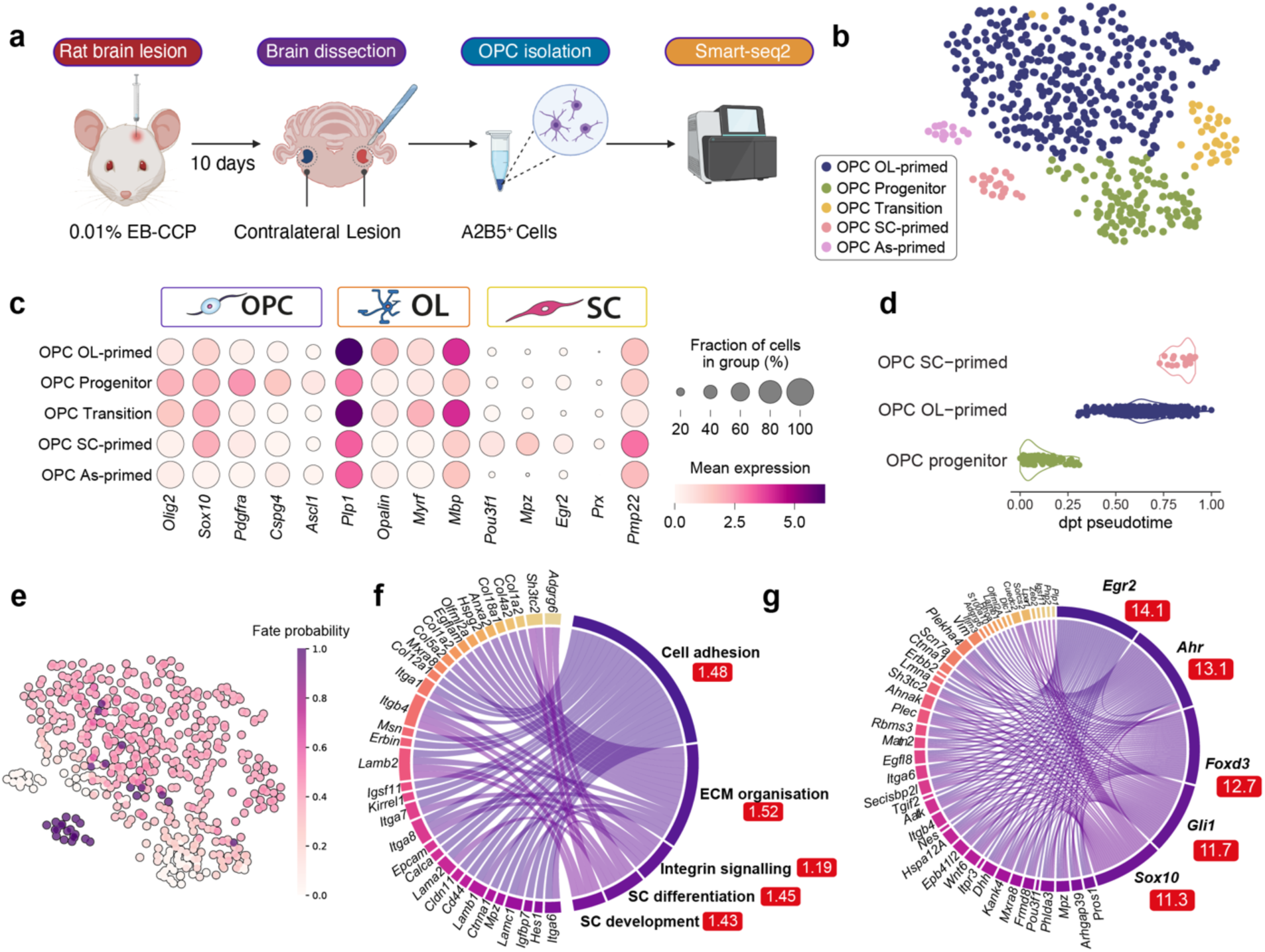
Single-cell transcriptomic analysis of purified A2B5+ OPCs. **a**, Purification of A2B5^+^ OPCs purification for single-cell RNA profiling via Smart-seq2. OPCs from the lesioned caudal cerebellar peduncle (CCP) and the uninjured contralateral side were collected and submitted for sequencing. A total of six animals were used: three animals underwent unilateral lesioning, from which both the lesioned and uninjured contralateral sides were sequenced, and three unlesioned animals served as naïve controls. **b,** Unsupervised clustering of single-cell transcriptional profiles from 517 A2B5^+^ cells. Five different states of OPCs were identified using Leiden clustering. Based on marker genes (shown in Fig. 1c and Extended Data Fig. 1e), these clusters were annotated as oligodendrocyte-primed (OL-primed, in dark blue), OPC progenitor in green, SC-primed (coral), transitioning cells (yellow), and astrocyte-primed (pink). **c,** Expression of lineage-enriched genes across different clusters used for classification of the cell states. An extended list is shown in Extended Data Fig. 1e, d. Diffusion pseudotime values across different cell clusters. Each dot represents a cell and the distribution is shown as a violin plot. **e,** Fate probabilities of individual cells transitioning toward a SC-like state. **f,** Pathway enrichment analysis of the genes significantly associated with the cell fate probabilities shown in Fig. 1e. The enrichment scores are shown in the red boxes. The top 200 genes were analysed on STRING, and only the top 5 pathways are shown. An extended table is available here: https://izu0421.github.io/opc2sc. **g,** Transcription factor analysis of the top 200 cell fate drivers. The enrichment scores are shown in the red boxes. Only the top 5 transcription factors from the GTEx library analysis of ChEA3 are shown. An extended table is available here: https://izu0421.github.io/opc2sc.

Unsupervised Leiden clustering identified five distinct clusters (Fig. 1b). We identified a SC-primed population significantly enriched in the lesion environment (Extended Data Fig. 1c), characterised by the upregulation of canonical SC markers (Extended Data Fig. 1d), —including (*Pou3f1, Mpz, Egr2, Prx,* and *Pmp22)*—alongside retained OPC transcripts. The remaining populations comprised a progenitor (*Pdgfra, Cspg4*), oligodendrocyte-primed (*Plp1, Myrf, Mbp*), transitioning, and astrocyte-primed (*Gfap, Aqp4*) clusters (Fig. 1c and Extended Data Fig. 1e)

To confirm the lineage commitment of these SC-primed cells, we performed trajectory analysis via graph abstraction and CellRank2 fate modeling. The SC-primed OPCs exhibited the highest pseudotime values and absorption probabilities toward a SC fate (Fig. 1e, and Extended Data Fig. 1f). Correlation of fate probabilities with gene expression identified extracellular matrix (ECM) organization and integrin signaling as the primary pathways driving this transition, with *Egr2* and *Sox10* pinpointed as key driver genes (Fig. 1f, g). We also identified the Wnt pathway previously implicated in the induction of SC differentiation by OPCs^14,17^ (Extended Data Fig. 1g).

### Identification of oSCs in active MS lesions from post-mortem human samples

To determine if this plasticity is a conserved feature of the human regenerative response, we applied a cell-type classifier, trained on our rat SmartSeq2 dataset, to 3,945 OPC nuclei extracted from a multiple sclerosis (MS) dataset (GSE279183^18^) (Fig. 2a, Extended Data Fig. 2a-d). This identified a discrete population of SC-primed OPCs (Fig. 2b). These nuclei are shared across three patients with chronic active lesions and are co-clustered with nuclei from patients with chronic active lesions (Fig. 2c, Extended Data Fig. 2b). The predicted oSCs also express SC-related genes including (*Sox10*, *Erbb3*, *Pmp22*, *Scn7a*). (Fig. 2d and Extended Data Fig. 2e).

**Fig. 2:**
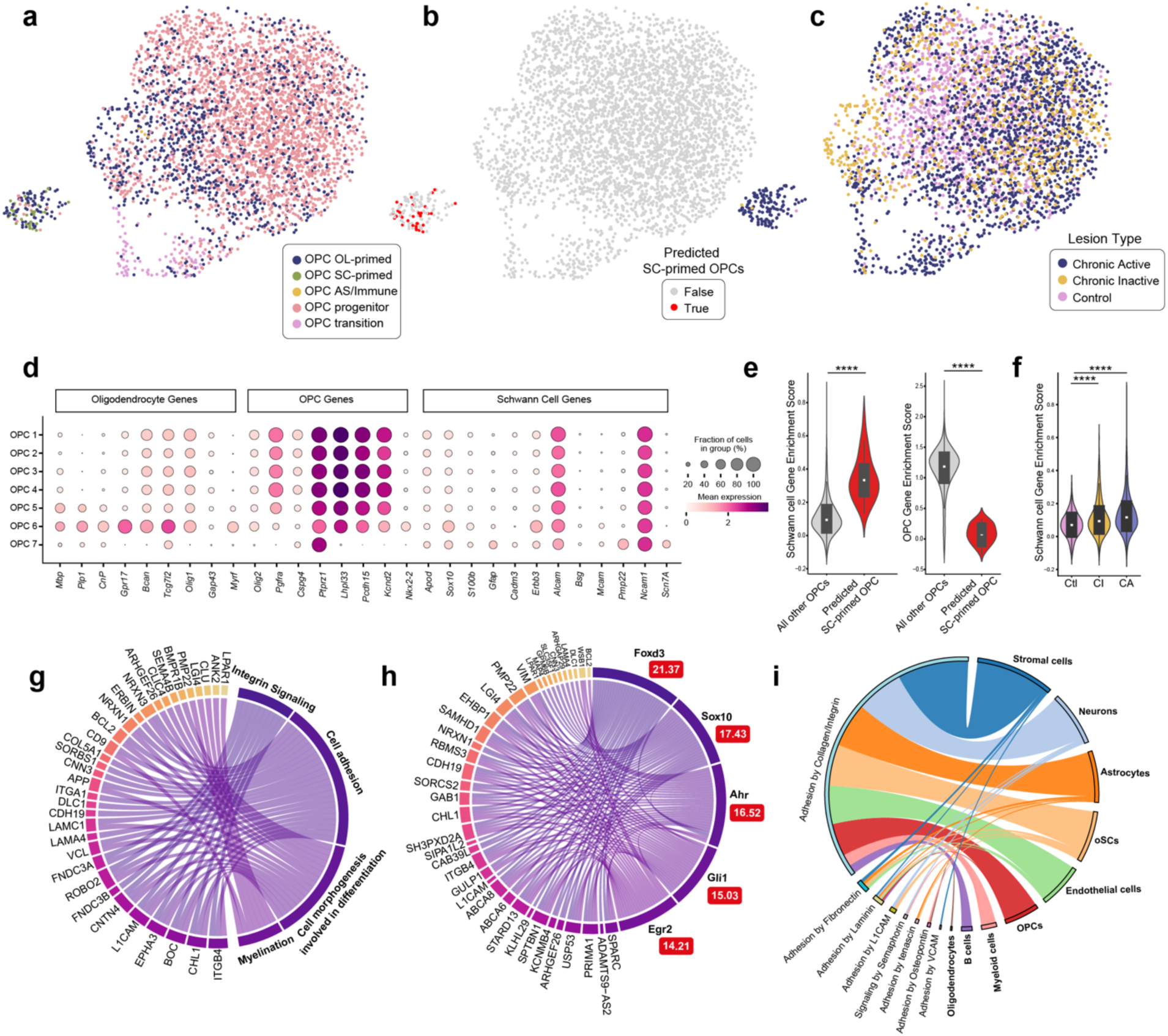
Identification of putative oSCs in OPCs from patients active MS lesions. **a**, Human OPC cell states from the Lerma-Martin et al., 2024 dataset were predicted using a CellTypist model trained on scRNA profiling of rat OPCs as described in Fig. 1. The human OPCs were mapped onto the embeddings from the rat Smartseq2 dataset to assign cell type identities. **b,** UMAP projection highlighting OPCs with a predicted SC-primed identity, coloured in red, while all other cells are shown in grey. **c,** UMAP projection of OPCs from the Lerma-Martin et al., 2024 dataset, with cells coloured by disease category. **d,** Expression of myelin genes, nerve support genes, and SC genes across the 7 Leiden clusters. These clusters were determined from independent clustering of the dataset and is further detailed in Fig. S2A. **e,** Expression of SC genes (left; ****p = 2.058 e-18) or OPC genes (right; ***p = 3.461e-21) in predicted oSCs compared to all other OPCs. Statistical significance was determined by a Mann-Whitney U test. **f,** Comparison of SC gene scores from OPCs isolated from control (Ctl), chronic inactive (CI) or chronic active (CA) human brain samples. Statistical significance was determined by a linear mixed effect model. **g,** The top 200 differentially expressed genes of the predicted oSCs compared to other OPCs and their associated biological pathways. **h,** The top 200 differentially expressed genes of the predicted oSCs compared to other OPCs and the predicted transcription factors that regulate them. The enrichment scores are shown in the red boxes. **i,** L-receptor interactions between oSCs and other cell types in the brain. The class of interaction by adhesion is on the left, and cell types on the right are highlighted in bold.

To further quantify the extent of lineage conversion, we calculated composite OPC and SC gene scores, based on well-established marker genes (Extended Data Table 1, Extended Data Fig. 2f). Predicted oSCs exhibited significantly higher SC and lower OPC gene scores compared to the rest of the OPC population (Fig. 2e). When stratified by lesion pathology, this SC-primed signature was significantly enriched in chronic active lesions compared to control tissue (Fig. 2f), whereas mature oligodendrocyte scores remained unchanged (Extended Data Fig. 2g).

Beyond gene markers, the cellular programs of these putative human MS oSCs mirrored our rodent findings, characterised by enriched integrin signaling (Fig. 2g) and the upregulation of identical core transcription factors (Fig. 2h). Interactome mapping using CellPhoneDB^19^ further identified a multifaceted signaling niche where stromal cells, neurons, astrocytes and oSCs themselves, provide the integrin-mediated cues necessary to facilitate this lineage switch (Fig. 2i and Extended Data Fig. 2h). While the presence of SCs in human pathologies has been documented for decades^5,6,8^, their origin has remained speculative. Our identification of this SC-primed OPC subpopulation provides the first transcriptomic evidence that these cells are not migrating in from the PNS, but are the progeny of a latent, neuroepithelial-to-neural crest plasticity inherent also to human progenitors.

### Vitronectin and BMP4 are sufficient to drive OPC to SC differentiation

Next, we aimed to functionally test our candidate pathways. While BMP signalling is upregulated within the injury niche, BMP alone directs OPCs toward an astrocytic rather than a SC fate^20,21^. This suggests within a lesion environment, other signals in addition to BMPs are required to generate SCs. Our transcriptomic analysis of SC-primed cells identified ECM-integrin engagement as a candidate for this missing signal. Among the ECM components upregulated during remyelination—including laminin α2, tenascins, osteopontin, and fibronectin^22,23^ — we identified vitronectin as the most likely mediator. Several molecules were excluded based on their temporal or spatial profiles: laminin α2 expression appears only after the onset of SC differentiation^22^, while tenascins are primarily astrocyte-derived and are associated with the suppression of SC differentiation from OPCs^24^. Additionally, our previous work showed that osteopontin deficient mice show no change in SC remyelination^23^, ruling it out as a required driver. By contrast, vitronectin, which is closely related to fibronectin, emerged as the primary candidate. It is a plasma-derived protein, and therefore present from early timepoints as the toxin mediated induction of lesions also causes the blood brain barrier to become leaky. Furthermore, vitronectin is sensed via the αV integrin expressed by OPCs^22,25–27^ and is reported to potentiate BMP signaling^28^. We therefore hypothesised that co-treatment with vitronectin and BMP4 might recapitulate the injury-specific signals^29–31^ responsible for directing OPCs toward a SC fate.

To test if these extrinsic signals are sufficient to trigger the SC program, we exposed primary OPCs to vitronectin and BMP4 (Fig. 3a). Within five days, OPCs grown under these conditions acquire a spindle-like morphology and formed wavelike patterns characteristic of primary SCs. These cells were immunoreactive to S100b, GFAP, and SOX10 with intensity and localisation identical to primary SCs (Fig. 3b). In contrast, adult OPCs on PDL-coated plates retained a round morphology and high OLIG2 and SOX10 expression (Fig. 3c). Single-factor treatments confirmed that neither vitronectin nor BMP4 alone is sufficient for oSC differentiation: BMP4 independently promotes astrogliogenesis, while vitronectin maintains progenitor identity but slightly reduces oligodendroglial differentiation potential (Extended Data Fig. 3). Thus, vitronectin serves as a second factor to redirect the outcome of BMP4 signalling away from astrocyte generation and towards a SC fate.

**Fig. 3:**
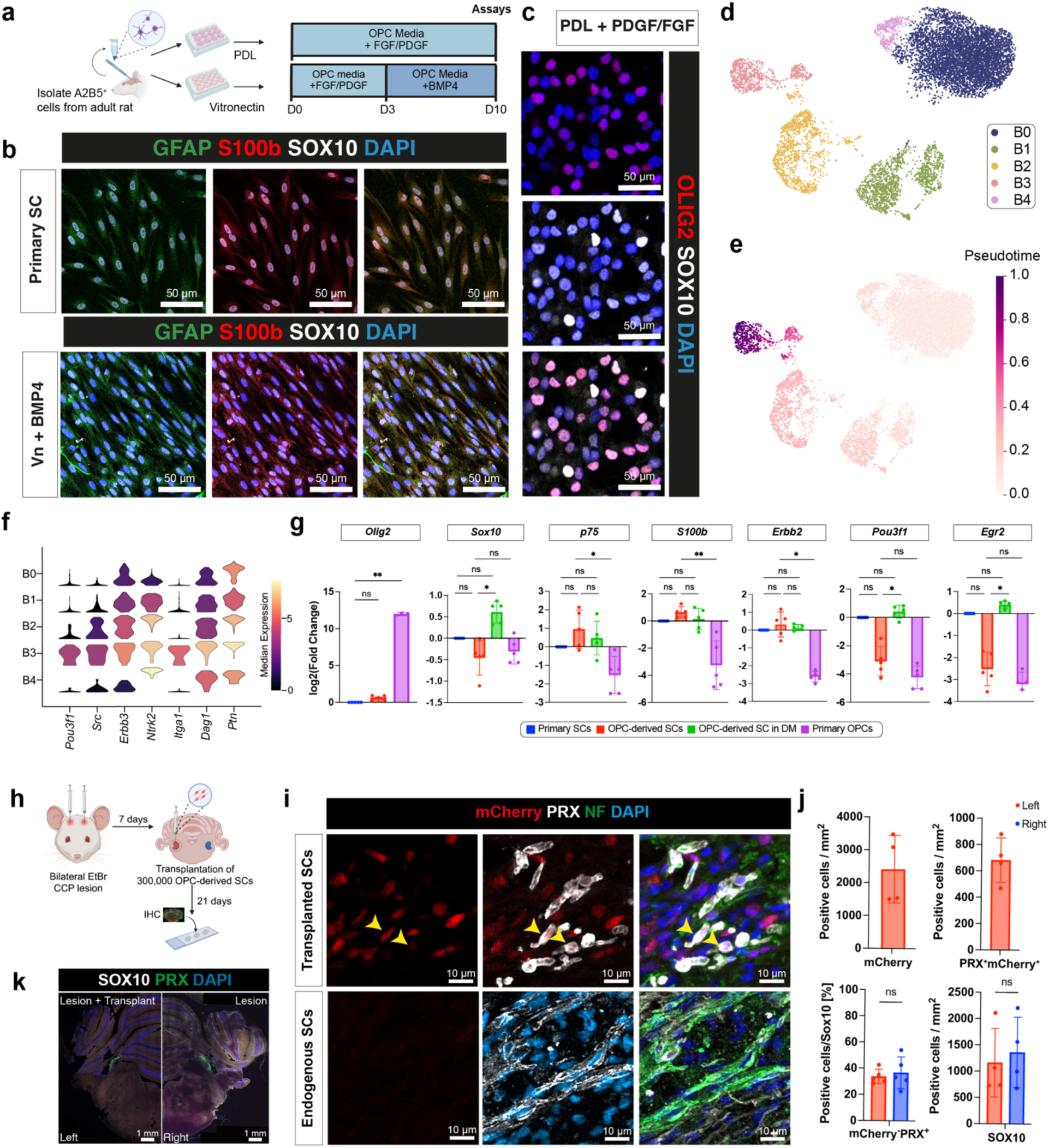
Vitronectin and BMP4 signalling is sufficient to generate transplantable, myelinating SCs. **a**, A2B5^+^ were isolated from the whole adult rat brain, then seeded on PDL or vitronectin coated plates. Control OPCs were plated on PDL and maintained in OPC media containing FGF/PDGF. To induce SC differentiation, OPCs were cultured on vitronectin (Vn), allowed to recover in standard OPC media for 3 days, then changed into BMP4-media for a week. **b,** GFAP, S100b and SOX10 immunostaining of cultured primary SCs and oSCs generated using Vn and BMP4. Data are representative of three independent experiments. **c,** OLIG2 and SOX10 immunostaining of OPCs cultured in OPC proliferation medium. Representative of three independent experiments **d,** UMAP representation of scRNA-seq data from Vn/BMP4-induced oSCs reveals five distinct cellular clusters. **e,** RNA velocity-predicted pseudotime of the Vn/BMP4-induced oSCs. **f,** Expression levels of SC genes in oSCs treated with Vn/BMP4, stratified by cluster. **g,** RT-qPCR of *Olig2, Sox10, p75, S100b, Erbb2, Pou3f1 and Egr2* expression in primary SCs, oSCs, oSCs in differentiation medium (DM) (containing cAMP and NRG1) and primary OPCs. Gene expression values are normalised to primary SCs. Values are mean ± SD. (*p< 0.05, **p< 0.01, by Kruskal-Wallis test with Dunn’s multiple comparison test; ns, not significant). Five biological replicates were used per group. **h,** Schematic overview of the brain lesioning and cell transplantation experiment. Bilateral EtBr brain lesioning was performed, then one week later 300,000 mCherry labelled oSCs were transplanted unilaterally into the left CCP. Animals were perfused 21 days after transplantation (28 days post-lesion, dpl) to assess remyelination. **i,** Immunohistochemistry of the transplanted area showing mCherryand PRX at 21 days post-transplant (28 dpl). Arrows indicate transplanted cells co-expressing mCherry and PRX. The bottom panel shows endogenous SC remyelination, identified by cells that are PRX^+^mCherry^-^. **j,** Quantification of the number of mCherry, SOX10 and PRX-positive cells within the lesion. **k,** Immunohistochemistry of brain lesioning shows SOX10, mCherry, and PRX protein expression.

To map the transcriptional programs driving the early stages of OPC-to-SC differentiation, we performed single-cell RNA sequencing using 10x Genomics on OPCs three days after treatment with vitronectin and BMP4. Unsupervised clustering resolved five cellular states referred to as B0 to B4 (Fig. 3d). Subsequent RNA velocity pseudotime^32^ revealed a single, continuous differentiation trajectory that originated from a progenitor-like state co-expressing initial SC genes (B1; *Pdgfra, Itga1, Dag1, Src)*, progressed through intermediate populations expressing SC genes such as *Pmp22* (B0, B4), and culminated in a single, terminal cluster (B3) that was most advanced in pseudotime (Fig. 3e). The direction of this trajectory was strongly correlated with the highest expression of SC markers (*Pou3f1, Src, Erbb3, Ntrk2, Itga1*) (Extended Data Fig. 4a). Despite this commitment, the B3 population exhibited a hybrid transcriptional profile: while highly enriched for SC-related genes, these cells had not yet completed the erasure of its oligodendroglial identity within this 3-day window, evidenced by the expression of *Olig1*, *Myrf*, *Opalin*, *Mag*, and *Mbp*, as well as co-expression of *Olig2* and *Sox10* (Fig. 3f and Extended Data Fig. 4b-d)

This finding led us to hypothesise that, like primary SCs, these immature oSCs would require additional extrinsic cues to complete their maturation into a myelinating cell. To test this, we transitioned newly induced oSCs into standard SC differentiation medium containing neuregulin and cAMP^33^. While newly induced oSCs expressed markers of immature SCs (*p75, S100b)* at levels comparable to primary SCs, their expression of pro-myelinating genes (*Pou3f1*, *Egr2*) remained low. In contrast, differentiated oSCs were transcriptionally indistinguishable from primary SCs across all markers tested (Fig. 3g). These findings demonstrate that vitronectin and BMP4 treatment initiates a switch to an immature SC fate, and that these oSCs are fully competent to respond to developmental cues, suggesting the potentially achieving a mature, myelinating phenotype.

### Extrinsic factor-induced oSCs exhibit stable lineage commitment and can myelinate CNS axons *in vivo*

Having established that adult OPCs can differentiate into SCs *in vitro* by exposure to a combination of vitronectin and BMP, we next asked whether SCs so generated could stably integrate and generate myelin sheaths around CNS axons using a focal model of ethidium bromide (EB)-induced demyelination. EB depletes oligodendrocytes and astrocytes, enabling remyelination by newly differentiated oligodendrocytes and SCs. We induced bilateral lesions, then mCherry-labelled oSCs were transplanted unilaterally into the left hemisphere 7 days later (Fig. 3h). This design establishes each animal as its own internal control, allowing us to directly compare the behavior of transplanted oSCs against the spontaneous regenerative response in an identical lesion environment. The lesioned area was assessed three weeks post-transplant (28 days post-lesion).

Since SC remyelination is a feature of spontaneous repair in EB-CCP lesions^34^, we expected total SC remyelination to involve contributions from both endogenous and transplanted SCs. We identified those that were of transplant origin by the co-expression of mCherry and PRX, a protein specific to myelinating SCs but not oligodendrocytes. No mCherry^+^ cells were detected in the non-transplanted contralateral lesions. The total amount of endogenous SC remyelination (quantified by the number of mCherry^-^PRX^+^ cells) in these control lesions remained consistent between the transplanted and the contralateral control lesion (Fig 3i-k). This indicated that transplanted oSCs successfully occupied and myelinated axons without displacing or inhibiting the differentiation of endogenous OPCs.

Furthermore, transplantation did not alter lesion size, microglia recruitment, or astrocytic hypertrophy between the left and right CCP (Extended Data Fig. 5), indicating that oSCs do not exacerbate the local cellular responses. This lack of reactive pathology is striking, as unlike transplanted peripheral-SCs, which are typically sequestered into isolated islands by exclusionary astrocytic responses^35^, transplanted Vn/BMP4-induced oSCs are stable and functional *in vivo* and show niche-compatible integration within the CNS.

### Injury-associated ECM signalling induces decreased expression of Olig2 but sustains levels of Sox10 in OPCs transitioning towards SCs

Having established niche-associated factors that will induce the differentiation of oSCs, we next identified the internal mechanisms governing this fate switch. The transcription factors SOX10 and OLIG2 are central to glial identity, but while they function synergistically to stabilise the oligodendroglial lineage, OLIG2 is considered a specific and essential pioneer factor for oligodendrocytes^36,37^, wherein SOX10 is also a master regulator for SC development^38^. We therefore hypothesised that OPC-to-SC differentiation requires the loss of OLIG2 to permit a SOX10-driven SC transcriptional program to override the default oligodendrocyte identity. Our rat single-cell data confirmed this signature: SC-primed OPC cluster expressed high levels of *Sox10*, but significantly lower levels of *Olig2* compared to the oligodendrocyte-primed cluster (Fig. 4a).

**Fig. 4:**
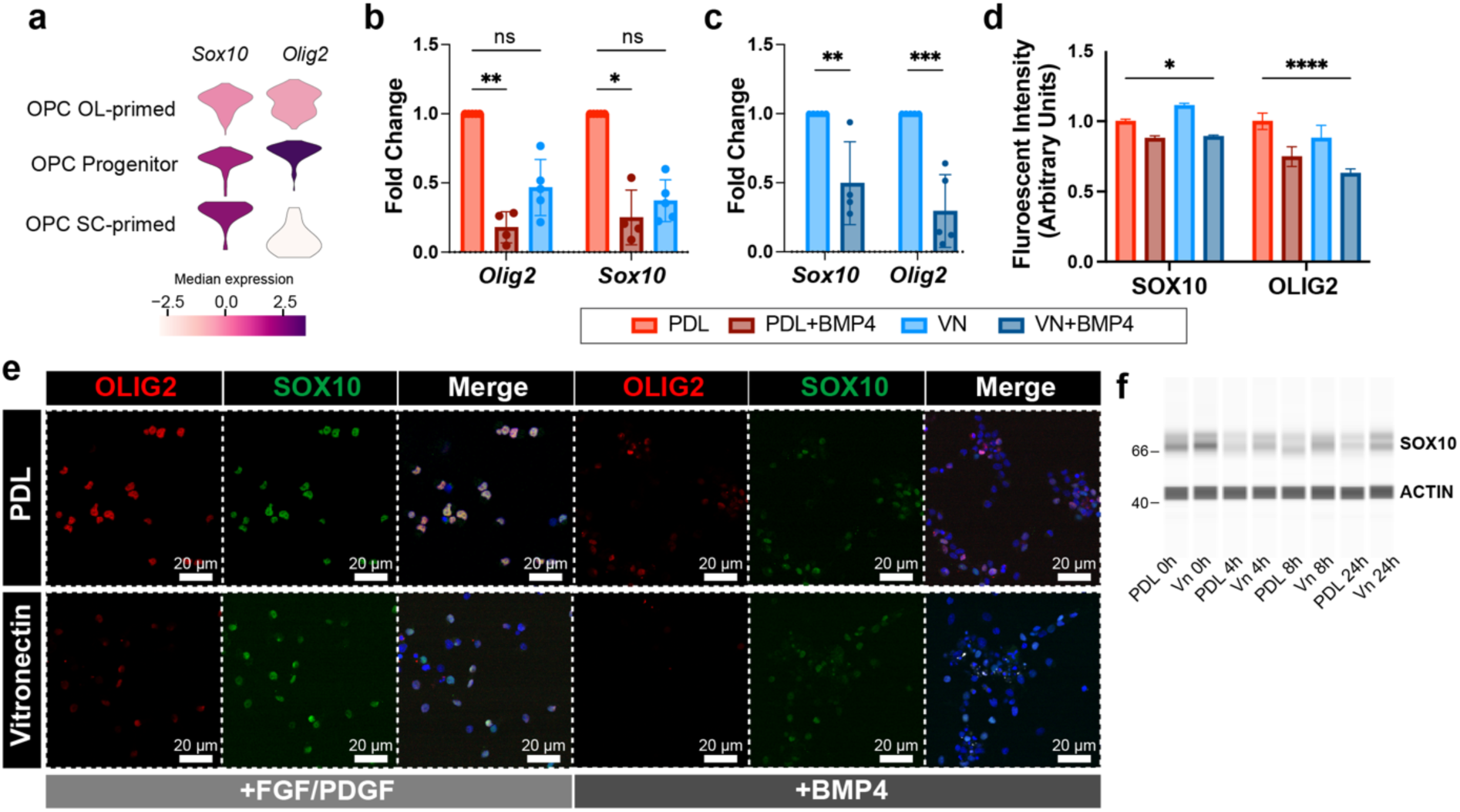
BMP4 decreases OLIG2 and SOX10 levels, but vitronectin stabilises SOX10 protein. **a**, Expression levels of *Olig2* and *Sox10* from the Smart-seq dataset, comparing SC-primed OPCs to OPCs remaining within the oligodendroglial lineage. **b,** RT-qPCR analysis of *Olig2* and *Sox10* mRNA expression in OPCs cultured on PDL or vitronectin, with or without BMP4 treatment. Values are mean ± SD, normalised to the PDL control condition. Statistical significance was determined by Kruskal-Wallis non-parametric test (*p < 0.05, **p<0.01), followed by Dunn’s post-hoc test. **c,** RT-qPCR analysis of *Olig2* and *Sox10* mRNA expression in OPCs cultured on Vn with or without BMP4 treatment. Data are normalised to samples cultured on vitronectin. Statistical significance was determined by Kruskal-Wallis non-parametric test (**p < 0.01, ***p<0.001), followed by Dunn’s post-hoc test. RT-qPCR experiments were conducted with four biological replicates per group. **d,** Quantification of fluorescent intensity of SOX10 and OLIG2 in OPCs cultured on PDL or vitronectin under standard proliferation conditions (+PDGF/FGF) or following BMP4 treatment. Values are normalised to cells cultured on PDL. Values are mean ± SD, by two-way ANOVA with Šidák’s multiple comparison test. **e,** Representative immunohistochemistry images of SOX10 and OLIG2 OPCs cultured on vitronectin or PDL with or without BMP4. Representative data from three independent experiments. **f,** OPCs were cultured on PDL or vitronectin (Vn) and treated with cycloheximide. A Jess Simple western blot shows SOX10 and ACTB protein levels at 0, 4, 8, and 24 hours. Experiment representative of three biological replicates.

To validate the predicted SOX10/OLIG2 switch, we assessed the effects of vitronectin and BMP4 on these core regulators. Vitronectin significantly reduced *Olig2* transcript and protein levels, an effect amplified by BMP4. Intriguingly, while *Sox10* mRNA similarly decreased, SOX10 protein was robustly maintained in the nucleus of cells treated with vitronectin and BMP4 (Fig. 4b-e). Cycloheximide chase assays confirmed a post-transcriptional mechanism: the half-life of SOX10 protein was significantly extended in the presence of vitronectin and BMP4 compared to controls (Fig. 4f).

### Direct SOX10/OLIG2 manipulation recapitulates niche-induced fate switching

We next addressed whether direct genetic manipulation of the SOX10/OLIG2 circuit is sufficient to bypass the extrinsic injury niche cues and drive the oSC fate switch. For this, we generated and validated lentiviral constructs to overexpress *Sox10* or to knock down *Olig2* levels using an *Olig2* shRNA lentivirus (Extended Data Fig. 6a-e). This direct genetic modulation was sufficient to phenocopy the effects of vitronectin and BMP4, with the cells undergoing a profound phenotypic conversion into spindle-like, wave-forming SCs that were transcriptionally and morphologically indistinguishable from primary SCs (Extended Data Fig. 6f). These cells demonstrated full functional competency: they matured in response to cAMP/neuregulin, myelinated DRG axons *in vitro*, and remained proliferative over multiple passages without loss of identity (Extended Data Fig. 7). Furthermore, comparative single-cell transcriptomics revealed that SOX10/OLIG2-induction and Vn/BMP4 treatment converge on an identical mature SC-like state. Trajectory analysis demonstrated that direct SOX10/OLIG2 manipulation achieved this fate with significantly higher efficiency than with extrinsic factors (Extended Data Fig. 8). These data establish that the SOX10/OLIG2 axis is the sufficient central integrator of the injury niche, providing a potent and deterministic driver of the oSC fate switch.

### AAV vector-mediated modulation of Olig2 and Sox10 can increase SC remyelination *in vivo*

To determine if the SOX10/OLIG2 circuit could drive repair *in situ*, we delivered a dual-promoter AAV vector to overexpress HA-tagged SOX10 and to knock out *Olig2* (hereafter referred to as AAV-SOX10/OLIG2) in OPCs (Fig. 5a, b). Following focal demyelination using lysolecithin^39,40^, this cell-autonomous manipulation bypassed niche requirements, forcing endogenous OPCs to adopt a myelinating SC fate. Indeed, AAV-SOX10/OLIG2 treated animals exhibited a twofold increase in peripheral-type myelin compared to AAV-mKate2 controls (Fig. 5c-g), with no significant differences in lesion size, microglial response, or overall astrogliosis (Extended Data Fig. 9a, b).

**Fig. 5.**
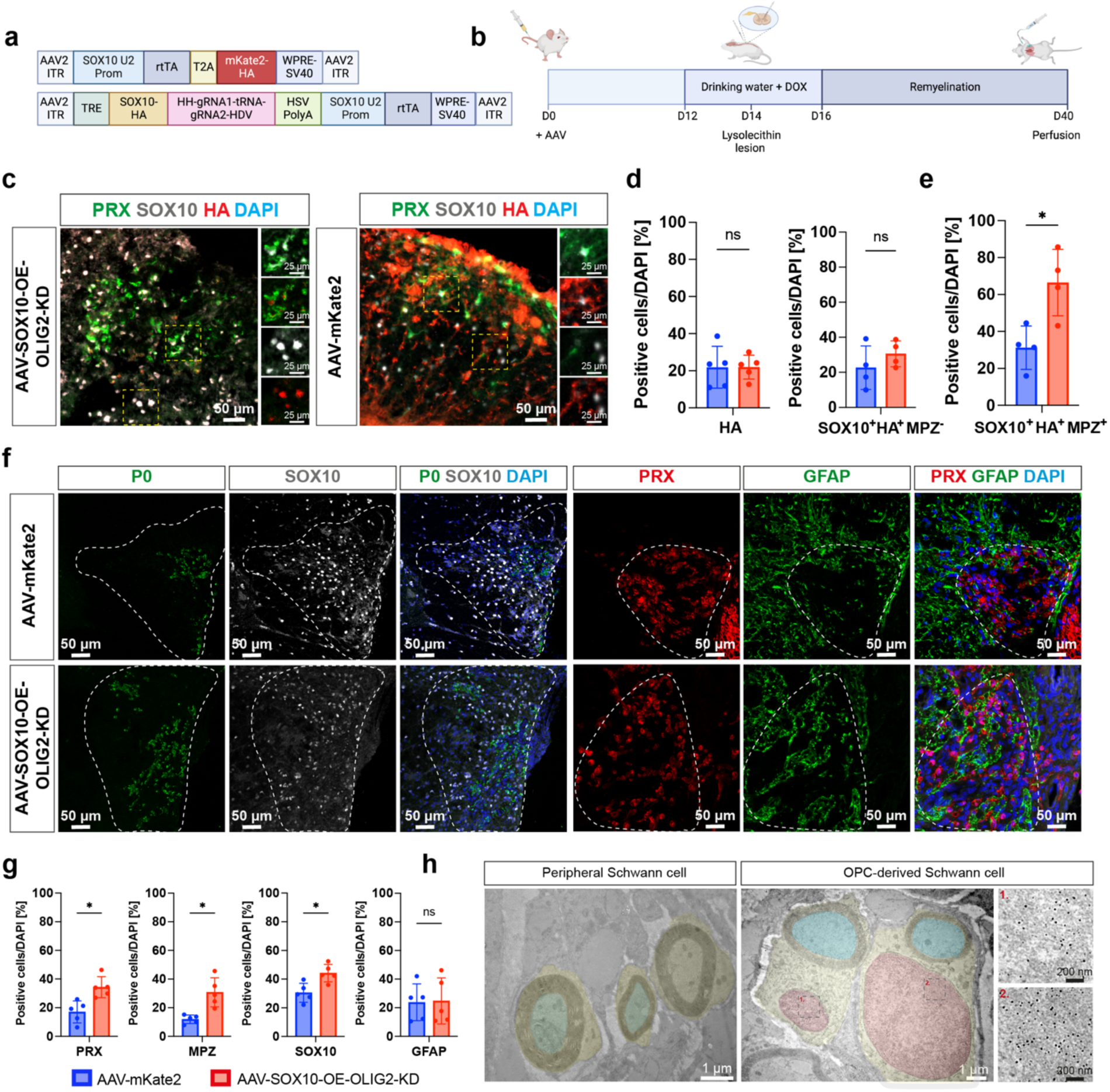
*Olig2* downregulation and *Sox10* overexpression are sufficient to induce OPC differentiation into SCs *in vivo*. **a**, Schematic linear map of the AAV-SOX10-OLIG2-gRNA and AAV-mKate2 plasmid constructs. **b,** Schematic of the experimental overview. Animals received either AAV-SOX10/OLIG2 or AAV-mKate2 via tail vein injection, followed by doxycycline administration in drinking water from D12 to D16. At D14 post-injection, a focal lysolecithin-induced demyelinating lesion was created in the spinal cord. Remyelination was assessed at D40, three weeks after demyelination. **c,** Representative immunohistochemistry images of lesioned spinal cords at 21 dpl showing SOX10^+^PRX^+^HA^-^ cells (endogenously arising SCs) and SOX10^+^PRX^+^HA^+^ cells (oSCs infected by the AAV) in both treatment groups. **d,** Quantification of HA+ and SOX10^+^ MPZ^+^HA^-^ cells in the spinal cord lesion. **e,** Quantification of SOX10^+^ MPZ^+^HA^+^ cells in the spinal cord lesion. **f,** Immunohistochemistry of the lesion area PRX, MPZ, and GFAP. **g,** Quantification of **(f)** shows a significant increase in the number of PRX*^+^*, MPZ*^+^*, and Sox10*^+^* cells in animals treated with AAV-SOX10/OLIG2 compared to AAV-mKate2-treated animals. Values are presented as mean ± SD. (*p< 0.05, ns, not significant, by unpaired t-test). The above data was obtained from four biological replicates per group, with at least three fields of view quantified per replicate. b, Immuno-electron microscopy image showing the ultrastructure of an HA^+^ SCs within the spinal cord. The SC body is pseudocoloured yellow, the associated axon in blue and the SC nucleus in red. The inset highlights immunogold particle labelling of HA in the SC nucleus.

Crucially, direct SOX10/OLIG2 manipulation enabled oSCs to override the exclusionary astrocyte-SC boundaries typically observed in CNS pathology^35^. While SCs in control lesions were restricted to astrocyte-free zones, in animals receiving AAV-SOX10/OLIG2, oSCs successfully integrated deep within GFAP^+^ territories (Fig. 5f and Extended Data Fig. 9c). This enhanced integration of SCs within astrocyte-rich lesion areas occurred without any corresponding change in the overall astrocyte response to demyelination between the control and AAV-treated groups. Immuno-electron microscopy for HA confirmed that these induced cells formed stable, compact peripheral-type myelin sheaths with a characteristic 1:1 axonal ratio and basal lamina (Fig. 5h). Thus, direct modulation of the SOX10/OLIG2 circuit expands the scale of the regenerative response and enables repair glia to navigate a hostile CNS environment.

## DISCUSSION

The spontaneous generation of neural-crest-derived Schwann cells (SCs) from neuroepithelial OPCs represents a rare instance of cross-germ layer differentiation during vertebrate regeneration. While earlier reports of adult stem cells crossing germ-layer boundaries were confounded by cell fusion^41^, the OPC-to-SC transition is a fate switch that defies traditional definitions of de-differentiation^42,43^ or transdifferentiation^44^. We find no evidence of OPCs reverting to a common neuroectodermal/neural crest progenitor, but instead it represents a form of adaptive cellular reprogramming triggered by injury^45^. Since OPCs are adult progenitors rather than terminally differentiated cells, this cross-germ layer transition represents a unique class of plasticity that extends beyond current paradigms of adaptive reprogramming.

This plasticity is striking given the early developmental divergence of the neural crest from the neuroepithelium at the time of gastrulation, occurring between embryonic days 6 and 9 in mice and at around 3 weeks in humans. This divergence is conserved back to the most primitive myelinated vertebrates (*Chondrichthyes*)^46^, highlighting the evolutionary separation of the central and peripheral nervous systems. Our findings demonstrate that this developmental boundary is not absolute, but rather a conditional state maintained by the environmental regulation of a transcriptional gatekeeper. By defining OPCs as a population capable of such transitions challenges the perceived rigidity of glial identity, adult stem cell differentiation potential, and expand the current paradigms of adaptive reprogramming.

We propose that this fate switch is governed by a dual-checkpoint control mechanism. Under homeostatic conditions, astrocyte-derived BMP antagonists like SOSTDC1 reinforce oligodendrocyte commitment^14,17^. Following injury, the loss of astrocytic inhibition creates a window of plasticity where unopposed BMP signaling destabilises the OLIG2-SOX10 complex and attenuates the oligodendroglial program. Notably, this disrupted state is metastable, as without further instruction BMP alone is insufficient for SC conversion, and OPCs default instead toward an astrocytic fate^20,21^. We identify vitronectin—a plasma-derived ECM protein that enters the parenchyma following blood-brain barrier compromise—as the critical second factor. By activating αV-family integrins, vitronectin signaling (likely via PI3K or JNK pathways) converges with BMP activity to stabilise SOX10^22^. In OPCs, this shifts transcriptional partnerships from the canonical OLIG2-SOX10 complex to a SOX10-POU3F1 interaction, effectively diverting OPCs from an oligodendroglia program to a SC fate. This dual-input requirement ensures that the cross-germ layer transition is spatially restricted to the specific environment of the injury niche and therefore explains why SCs are typically not observed in the homeostatic CNS.

Our identification of a discrete SC-primed OPC population in human MS lesions establishes this plasticity as a conserved feature of human disease. These rare progenitors are characterised by the expression of a SC signature alongside the incomplete erasure of oligodendroglial identity. The finding that AAV-mediated SOX10/OLIG2 manipulation significantly increased the volume of PRX^+^/P0^+^ remyelination *in vivo* (Fig. 5e) demonstrates that there is scope to further increase the contribution of SC remyelination as a therapeutic strategy for demyelinating diseases.

While the evolutionary drivers of SC-mediated CNS repair remain elusive, its translational potential is profound. In MS, peripheral-type myelin produced by SCs may provide an “immune-privileged” form of repair, inherently resistant to the autoimmune targeting of CNS epitopes. Furthermore, the rapid kinetics of SC-mediated remyelination^47^ and their robust support for axonal regeneration^48^ —in contrast to the inhibitory environment of NOGO- or MAG-expressing CNS myelin^49,50^ - could significantly reduce the window of axonal vulnerability following injury. Historically, the mutual exclusivity of SCs and reactive astrocytes has limited the scope of such repair^51,52^. However, our demonstration that targeting the SOX10/OLIG2 axis can bypass these inhibitory signals to enable astrocyte-compatible SC remyelination offers a transformative strategy for biasing OPC fate choices in chronic CNS pathology.

## Acknowledgements and Funding

C.Z.C. was supported by a Wellcome Trust PhD fellowship. N.M. was supported by the Sackler Trust. B.N. was supported by a Kim and Julianna Silverman research fellowship. K.S.R was supported by a Banting Postdoctoral Fellowship from the Canadian Institute of Health Research as well as a Multiple Sclerosis Society of Canada Fellowship. R.J.M.F. is supported by funding from the UK Multiple Sclerosis Society (MS50) and Dr Miriam and Sheldon G Adelson Medical Research Foundation, and a core support grant from the Wellcome Trust and MRC to the Wellcome Trust-Medical Research Council Cambridge Stem Cell Institute (203151/Z/16/Z). We thank Gemma Cronshaw for performing the tail vein injections, Maike Paramour and Laura Everton for generating the 10X sequencing libraries. We thank Karin Müller and Louis Elfari (University of Cambridge Stem Cell Institute, for their assistance with the immunoelectron microscopy.

## Author Contributions

Conceptualisation, C.Z.C., B.N., R.J.M.F.; Methodology, C.Z.C., N.M., Y.Y, B.N.; Investigation and data analysis, C.Z.C., N.M., Y.Y., J.F.C., K.S.R., C.Z., M.H., B.N.; Writing C.Z.C., B.N., R.J.M.F.; Funding Acquisition R.J.M.F.; Supervision, P.A-F., B.N., R.J.M.F.

## Data, code and materials Availability

All code is available in our GitHub repository (https://izu0421.github.io/opc2sc). All single-cell RNA sequencing data are publicly available in the GEO repository. The Smart-Seq2 rat lesion OPC data are accessible under accession number GSE311740. The 10x Genomics *in vitro* OPC and SC data are accessible under accession number GSE312112.

## Materials and Methods

### Animal Husbandry

All procedures were performed in compliance with the rules and regulations of the UK Home Office and were approved by the University of Cambridge Animal Welfare and Ethical Review Body. Animals were housed in groups of up to 4 animals and kept at a 12-hour dark-light cycle with free access to standard chow and water. Animals were randomly allocated to experimental groups, which were matched for age and gender.

### Isolation and culture of oligodendrocyte progenitor cells

Primary OPCs were isolated from neonatal (P2-P10) or young adult (8-24 week) male and female Sprague Dawley rats. The brains were diced with a scalpel into 1 mm^3^ pieces. The tissue pieces were digested in Hibernate A Low Fluorescence (HALF, #HALF500) solution containing 34 U/mL papain (Worthington, #LS003127) and 40 μg/mL DNase I (Sigma, #D5025) on an orbital shaker (55 rpm) for 40 min at 37°C. The tissue was centrifuged then resuspended in HALF supplemented with B27 (1:50; Gibco #17504044) and sodium pyruvate (2 mM; Gibco #11360070). To obtain a single cell suspension, the tissue was triturated 10 times each using a 5 mL serological stripette, and 3 fire-polished pipettes of decreasing diameters. The supernatant, containing the cells, was filtered through a 70 μm cell strainer into a tube containing 22.5% isotonic Percoll (GE Healthcare, #17-0891-01) in DMEM/F12 (Gibco, #31331028). The single cell suspension was then separated from tissue and myelin debris by gradient density centrifugation at 800 g (without brakes) for 20 min at RT. The resulting cell pellet was resuspended in red blood cell lysis buffer (1 mL, BD Biosciences, #555899) for 1 min to remove contaminating red blood cells. Subsequently, the cell suspension was incubated in Miltenyi washing buffer (MWB; 2 mM EDTA, 2 mM sodium pyruvate, 0.5% BSA) and 1.7 μL mouse-anti-rat-A2B5-IgM antibody (Merck Millipore, #MAB312) for each adult brain, or 0.25 μL antibody for each neonate brain for 15 min at 4°C. After incubation, cells were washed with 7.5 mL MWB and centrifuged (300 g, 5 min, RT). The pellet was resuspended in 80 μL MWB-insulin, containing 20 μL rat anti-mouse IgM magnetic beads (Miltenyi, #130-047-302) for 15 min at 4°C. The secondary antibody was diluted in 7 mL MWB buffer and centrifuged (300 g, 5 min, RT). After the spin, cells were resuspended in 0.5 mL MWB-insulin, then transferred into the MS-column. The column was washed three times with 0.5 mL MWB-insulin per wash, then discarded. To collect the A2B5^+^ bound fraction, the MACS MS column was removed from the magnetic stand, and cells were eluted with 1 mL OPC medium (60 μg/mL N-Acetyl cysteine (Sigma, #A7250), 10 μg/mL human recombinant insulin (Gibco, #12585014), 1 mM sodium pyruvate (Gibco, #11360070), 50 μg/mL apo-transferrin (Sigma, #T1147), 16.1 μg/mL putrescine (Sigma, #51799), 40 ng/mL sodium selenite (Sigma, #214485), 60 ng/mL progesterone (Sigma; P0130), 330 μg/mL bovine serum albumin (Sigma, #A4919). Cells were counted, then 20,000 cells/cm^2^ were plated onto poly-D-lysine (5 μg/mL PDL; Sigma)-coated glass bottom well plates and incubated in OPC medium supplemented with b-FGF (20 ng/mL; Peprotech, #100-18B) and PDGF (20 ng/mL; Peprotech, #100-13a) at 37°C, 5% CO_2_, 5% O_2_. Counts averaged 1.2 x10^6^ OPCs per brain. The morning after isolation, the OPC proliferation medium was completely replaced to remove any dead cells. Whilst in culture, two-thirds of the media was replaced every 48 h with growth factors added freshly per media change. Cells were allowed to recover for up to 72 h after isolation in proliferation media before treatment with small molecules, lipofectamine transfection or viral infection.

### Isolation and culture of Schwann cells

SCs were obtained from the sciatic nerve of 1-2 day old Sprague Dawley rat pups. Briefly, sciatic nerves were dissected and immediately submerged in HALF isolation medium. The epineurium was removed, and the peeled nerves were teased until the fascicles were separated and cloudy. The nerves were transferred to a fresh tube containing cold HALF solution and collected by centrifuging for 5 min at 300 g at RT. The tissue was then digested in collagenase type 1 (2 mg/mL; Sigma, #C0130) and trypsin (1.25 mg/mL, Sigma, #T9201) diluted in HALF at 37°C for 45 min. To obtain a single cell suspension, the nerves were mechanically dissociated by triturating with a P1000 pipette. Cells were centrifuged (3 min, 300 g, RT), and then seeded onto laminin (Sigma, #L2020)-coated well plates with fresh D-media (DMEM media supplemented with 10% FBS) and left to recover in the incubator overnight (37°C, 5% CO_2_). The following day, cultures were treated with cytosine arabinoside (Ara-C; 10 μM; Sigma, #C6645) diluted in D-media for 2 days to suppress growth of fibroblasts. Subsequently, the Ara-C containing media was removed, and the cells were allowed to recover and expand for 2 days in normal D-media. The remaining fibroblasts were eliminated by complement-mediated killing using the IgM class anti-Thy1.1 antibody (1:500; Serotec, #MCA04G) and rabbit complement (Millipore, #234400). To promote SC expansion, the cultures were switched onto SC growth media containing DMEM supplemented with 10% FBS, neuregulin (10 ng/mL, Peprotech, #396-HB), forskolin (2 μM; Sigma, #F6886), and 1% penicillin/streptomycin. Cells were cultured at 37°C, 5% CO2 and fed every 3 days. After reaching 80% to 90% confluency, SCs were trypsinised with 0.25% Trypsin-EDTA, centrifuged (5 min, at 300g, RT) and passaged.

### Isolation and culture of dorsal root ganglion (DRG) cells

DRGs were collected from the spinal columns of E14 rat pups and transferred into a 6 cm sterile dish containing 1 mL ice-cold HALF. Then the DRGs were transferred into a falcon tube containing 2.5 mL 0.025% trypsin dissolved in Ca^2+^ and Mg^2+^-free PBS (Sigma, #T9201) and incubated at 37°C for 20 min on an orbital shaker. Afterwards, the ganglia were further digested in 600 μL type I collagenase solution (1.5 mg/ml; Sigma, #SCR103) at 37°C for 30 min on an orbital shaker. Subsequently, DMEM with 10% FBS was added to terminate the digestion, and the cell pellet was collected by centrifuging for 5 min at 300 g at RT. To obtain a single cell suspension, the cell pellet was mechanically dissociated by triturating with a P1000 pipette. Cells were centrifuged (3 min, 300 g) and then seeded onto laminin coated well plates with serum free DMEM (DRG media) containing 50 ng/mL NGF (R&D systems, #256-GF), N2 supplement (1:50, Gibco, #17502048) and 0.05% BSA, and left to recover in the incubator overnight (37°C, 5% CO_2_). The following day, cultures were treated with Ara-C (10 µM; Sigma) diluted in fresh media for 2 days to suppress growth of fibroblasts and SCs. Subsequently, the Ara-C containing media was removed, and the cells were allowed to recover and expand for 2 days in normal media. Cultures were cycled through Ara-C mediated elimination of nonneuronal cells at least three times to obtain purified neuronal cultures.

Once the DRG neurons were purified, SCs, prelabelled with a PGK-H2B-Cherry (Addgene plasmid #21217) lentivirus, were trypsinised, then seeded with DRGs at a density of 2 × 10^5^ cells/400 µL. DRG-SC co-cultures were maintained in DMEM (Gibco, #D5796) supplemented with 10% FBS, Glutamax (2mM, Gibco), NGF (50 ng/mL NGF) and penicillin-streptomycin (C media). After 7 days, ascorbic acid (50 µg/mL) was added to the C medium to induce myelination. Cells were maintained at 37°C, 5% CO_2_ with media changes every 2 to 3 days.

### shRNA-lentivirus mediated deletion of Olig2

Oligonucleotides encoding the Olig2 shRNA sequence (Fwd: GATCCGTTCTTGTGTGTGTAAATATAACTCGAGTTATATTTACACACACAAGAACTTTTT; Rev: AATTCAAAAAGTTCTTGTGTGTGTAAATAT AACTCGAGTTATATTACACACACAAGAACG) were oligoannealed, then ligated into the pGreenPuro-shRNA-Stx3S-C4 (Addgene #99742) using a T4 ligase (New England Biolabs). The sequences were confirmed with Sanger sequencing, then plasmids were transformed into Stable Competent E.coli (New England Biolabs, #C3040H) and purified with the NucleoBond Xtra MIDI kit (Machery-Nagel, #11932492).

To generate shRNA lentiviruses, HEK293T cells were grown in 15-mm plastic dishes and triple-transfected with the pMD2.G envelope plasmid (Addgene plasmid #12259), the psPAX2 packaging plasmid (Addgene plasmid #12259), and the transgene plasmid. Viral supernatant was collected at D3 and D5 post transfection, then concentrated using the Lenti-X Concentrator (TakaraBio, #631232) as per manufacturer’s instructions. Purified lentiviruses were aliquoted and stored at -80°C until required. To knockdown *Olig2*, OPC cultures were treated with purified shRNA-lentivirus diluted in 500 μL of OPC media containing growth factors (PDGF and FGF) for 48 h.

### Construction of gRNA and overexpression plasmids

Two AAV plasmids were generated using PCR to amplify fragments containing appropriate overlaps, followed by Gibson assembly using the NEBuilder® HiFi DNA Assembly Master Mix (NEB, #E2621S). Detailed plasmid maps and cloning primers are provided in the supplementary GenBank files.

#### 1. AAV-TRE-HA-Sox10-gRNA-Sox10U2PROM-rtTA-WPRE

This construct was assembled from elements derived from several templates, including: an AAV backbone (from AAV-CMVc-Cas9, Addgene #106431), a TRE promoter (from LV-pInducer20, Addgene #44012), and an HA-tagged Sox10 coding sequence (from Tet-O-FUW-Sox10, Addgene #45843). It also includes a synthetic gRNA expression cassette containing the first 110 bp of the mouse MALAT1 triple helix^53^, the Hammerhead and HDV ribozymes^54^, two *Olig2* targeting gRNAs (Olig2_gRNA1: ACGATGGGCGACTAGACACC, Olig2_gRNA2: GCACGACCTCAACATCGCCA) separated by a pre-tRNA-Gly sequence and a polyadenylation signal. The Sox10 U2 promoter (from LV-Sox10MCS5-GFP, Addgene #115783) and an rtTA-WPRE module (from pHR-EF1α-Tet-on 3G, Addgene #118592) were incorporated to enable doxycycline-dependent HA-Sox10 induction.

#### 2. AAV-Sox10U2PROM-rtTA-T2A-mKate2-WPRE

A second construct was generated using the same AAV backbone (AAV-CMVc-Cas9, Addgene #106431) and Sox10 U2 promoter, combined with rtTA, a T2A self-cleaving peptide, and WPRE elements (from pHR-EF1α-Tet-on 3G), together with mKate2 (from NFKBRp-mKate2-2xNLS-p2a-puroR, Addgene #82024) to enable fluorescent reporting of promoter activity.

### AAV Production

AAVs were produced and purified as described^55^. Briefly, HEK293T cells were grown in 15-mm plastic dishes and triple-transfected with the pHelper plasmid (Addgene plasmid #112867), the PHP-EB capsid plasmid (Addgene plasmid #103005), and the transfer plasmid. Virus particles were isolated by ultra-centrifugation (350,000 g) of the supernatant collected at D3 and D5 post transfection, as well as lysed cells collected at D5 post transfection using an Optiprep density-gradient medium (Sigma; #D1556). Following ultra-centrifugation, the viral layer was extracted, then concentrated using Amicon Ultra-15 centrifugal filter units (Sigma; #Z648043-24EA). The AAV titre was determined using SYBR Green qPCR and stored at 4°C until use.

### Induction of white matter lesions and Schwann cell transplantation in the brain

Adult female Sprague Dawley rats (12-20 weeks) were anesthetised with buprenorphine (0.03 mg/kg, s.c.) and 2.5% isoflurane. Then, bilateral demyelination was induced by stereotaxic injection of 4 μL of 0.01% ethidium bromide (EB) into the caudal cerebellar peduncles (CCPs), as previously described ^12^. The EB was delivered at a rate of 1 μL/min. 7 days later, 300,000 SCs pre-labelled with PGK-H2B-mCherry lentivirus (Addgene plasmid #21217) in 4 μL of HBSS^-/-^ were transplanted unilaterally into the ethidium bromide-induced area of demyelination in the left CCP.

### AAV delivery and spinal cord lesioning

5 X 10^11^ viral genomes of each AAV were tail vein injected into adult (8-12 weeks) ROSA-26-Cas9 (The Jackson Laboratory, #026179). To induce transgene activation, animals were switched onto water containing Ribena and doxycycline (2 mg/mL, Sigma, #D9891) from D10 to D14. Animals also received a one-time intraperitoneal injection of doxycycline on D11 (5 mg/ml, diluted in saline solution). 10 days after AAV delivery, spinal cord lesions were induced by injecting 1 μL of 1% lysolecithin (Sigma, #L4129) between two vertebrae of the thoracic column into the ventral funiculus at a rate of 1 μL per minute as previously described^34^. After lysolecithin delivery, the needle remained in position for an additional minute to prevent backflow of the lysolecithin solution.

### Perfusion and tissue processing for immunohistochemistry

Adult mice or rats were transcardially perfused with 0.9% NaCl solution for 1 minute, followed by 4% freshly depolymerised paraformaldehyde (PFA) in PBS for 7 minutes. The spinal cord and brain were dissected out then post-fixed in 4% PFA at 4°C overnight. Tissues were cryoprotected in 30% sucrose solution overnight, embedded in OCT medium (TissueTek, #4583), then cut into 12 μm thick sections using a cryostat.

### Immunohistochemistry of tissue slides or cultured cells

Non-specific antibody interactions were blocked by incubating tissue slides in blocking solution containing 10% donkey serum (Sigma, #D9663), 1% bovine serum albumin, 0.1% cold fish gelatin (Sigma, #G7041), 0.1% Triton X-100 (Sigma, #T8787) and 0.05% Tween20 (Sigma, #P9416) in PBS for 30 to 60 min at RT.

Primary antibodies (Extended Data Table 2) were diluted in 5% donkey serum in PBS and incubated overnight at 4°C. The slides were washed 3 times for 5 min with PBS, then incubated in secondary antibodies (Extended Data Table 2) diluted in 2% donkey serum in PBS (1 h, RT). Subsequently, nuclei were counterstained with DAPI (1:10,000) in PBS for 10 min, then washed 2 times with PBS for 5 min. The slides were mounted onto coverslips using Fluromount mounting media (Sigma, #F4680).

Similarly, cultured cells were washed with PBS containing Ca^2+^ and Mg^2+^ twice before fixation with 4% PFA for 15 min at RT. Subsequently, the cells were washed with 1X PBS three times (5 min, RT, shaking), blocked and permeabilised with 10% PBS-T (with 0.1% Triton X-100) for 30 min, then stained using the same protocol. Images were acquired using a confocal microscope (Leica TCS SP8, Zeiss LSM 980 Airyscan, or Zeiss Axioscan 7) and processed using ImageJ or QuPath. Cell populations were quantified manually from at least 3 fields of view per biological replicate.

### Perfusion and tissue processing for immunoelectron microscopy

Adult mice or rats were perfused transcardially with 0.9% NaCl solution for 1 minute, followed by a 6-minute perfusion of 4% freshly depolymerised PFA in PBS. The spinal cord was carefully dissected and post-fixed overnight at 4°C in a solution containing 2% PFA and 2% glutaraldehyde in PBS. After fixation, tissues were embedded in 10% gelatin (Sigma; #48724) and sectioned into 1 mm thick slices using a vibratome. These sections were further post-fixed in 4% PFA for 15 minutes, then blocked with a blocking buffer for 24 hours at 4°C.

Next, the tissues were incubated with the HA primary antibody (Invitrogen, #26183) at 4°C for 96 h. Following primary antibody incubation, the sections were treated with an immunogold secondary antibody (1:20; Sigma; G7652) for 24 h at 4°C. After staining, the immunogold signal was further developed using a silver enhancer kit (Sigma, #SE100) as per the manufacturer’s instructions. Stained sections were sections osmicated overnight at 4°C in a 0.25% osmium solution then dehydrated in an ascending ethanol series and propylene oxide then embedded in resin (TAAB). For electron microscopy analysis, ultrathin sections (70 nm) from the lesion sites were cut and transferred onto copper grids. These sections were stained with lead citrate, and grids were visualised using a Hitachi-HT7800 transmission electron microscope.

### RNA isolation and qRT-PCR

RNA was isolated from cultured OPCs using the Directzol RNA MicroPrep Kit (Zymo, #R2050). cDNA was generated using the QuantiTect Reverse Transcription Kit (Qiagen, #205311). For quantitative real-time (qRT)-PCR, PowerUp SYBR Green master mix (Applied Biosystems; #A25742), primers (500 mM, Extended Data Table 3), RNAse-free water, and cDNA were loaded according to manufacturer’s instructions. Each reaction was repeated in duplicate. Subsequently, RT-qPCR and melting curve analysis was performed on Life Technologies Quantstudio 6 Flex Real-Time PCR System. Fold changes in gene expression were calculated using the Livak method^56^.

### Western blot

Cells were lysed in RIPA lysis buffer (Sigma, #R0278) supplemented with 1% Halt protease and phosphatase inhibitor (Thermofisher, #78442) for 30 min on ice. The lysates were centrifuged for 30 min at 4°C at 13,000 g in a tabletop centrifuge. The supernatant was stored at 80°C. Protein quantification was carried out using the Pierce BCA protein assay kit (Thermofisher, #23209) measured with the SPECTROstar Nano absorbance plate reader.

For normal western blot, protein lysate was mixed with 4X NuPage LDS Sample Buffer (Thermofisher, #NP0007) and 10X NuPage Sample Reducing Agent (Thermofisher, #NP0009), then incubated at 70°C for 10 min. 50 μg of each protein sample were loaded per well, then separated on Bolt 4%–12% Bis-Tris Plus gels (Thermofisher, #NP0322PK2) in Bolt MOPS SDS running buffer (Thermo Scientific, #B000102) supplemented with 1 ml NuPage antioxidant (Thermofisher, #NP0005) at 150 V for 60 min. Subsequently, the protein was transferred to a polyvinylidene fluoride (PVDF, Immobilon, #IPFL00010) membrane in NuPage transfer buffer (Thermofisher, #NP0006) for 60 min at 20 V using the Mini Blot module (Thermo Scientific). After protein transfer, membranes were blocked in Intercept TBS blocking buffer (Li-COR, #927-60001) for 1 h at RT, then incubated in blocking buffer containing 0.1% Tween20 (TBS-T) containing primary antibodies on a shaker overnight at 4°C. The membranes were washed thrice in TBS-T for 5 min, and then incubated in secondary antibody diluted in blocking buffer, at RT for 1 h. After incubation, the membranes were re-washed three times in TBS-T and imaged using the Odyssey (Li-Cor).

For Jess Simple Western, protein lysates were diluted to a final concentration of 0.5 mg/mL, then all samples, and buffers were loaded into the cartridges as according to the manufacturers’ instructions. SOX10 (Abcam, #AB155279) was used at a concentration of 1:50, and bActin (Cell Signalling, #3700) was used at a concentration of 1:250 in Antibody Diluent 2 (Bio-Techne, #042-203).

#### Generation of single-cell RNA sequencing data for Smart-seq2

Focal white matter demyelination was induced unilaterally in the caudal cerebellar peduncle of adult female rats (12 weeks) using 0.01% ethidium bromide. 10 days post lesion, both the lesioned and contralateral uninjured CCP were microdissected, and A2B5^+^ cells were isolated using FACs. This timepoint corresponds to the onset of OPC differentiation in the EB-CCP lesion^34^. Sequencing libraries were prepared using a modified Smart-seq2 protocol^57^ and sequenced using the Illumina HiSeq 4000. Reads were aligned to the rat genome (Rattus_norvegicus Rnor_6.0.88).

### Generation of single-cell RNA sequencing data for 10X genomics

SOX10/OLIG2-induced oSCs and Vn/BMP4-induced oSCs were detached from culture plates using TrypLE Express (Gibco, #12604013) then 10,000 cells per sample were submitted to the Cambridge Stem Cell Institute Genomics core for RNA isolation and library preparation. Individual cells were partitioned into Gel Bead-In-Emulsions (GEMs) in accordance with the Chromium Single Cell 3’ protocol (10X Genomics, #120237). Briefly, the single cell GEMs were dissolved, which releases and allows mixing of the cell lysate with a primer pool containing an Illumina Read 1 sequence, a 16 nt 10x Barcode, a 10 nt Unique molecular identifier, and a poly-dT primer sequences). Incubation in this reverse transcription master mix yields barcoded, full-length cDNA. The GEMs were lysed, PCR amplified, and the quality of the cDNA was assessed using high sensitivity DNA electrophoresis (Agilent Technologies, #5067-4626) with the Agilent bioanalyzer system. Illumina-compatible cDNA libraries were prepared for sequencing using single cell the Single Cell 3’ Reagent kit protocol. Sequencing was performed on the Illumina NovaSeq 6000 S2. The number of assigned reads for each sample was between 362 million and 666 million, with a minimum read depth of 40,955 reads per cell. After sequencing, FASTQ files were mapped to the rat reference genome (RGSC6.0/rn6) using Cellranger v5.0.1 and processed on Scanpy.

### Analysis of single-cell RNA sequencing data

The analysis of single-cell RNA sequencing data was performed similarly to previous work^58^. For the *in vivo* data, low-quality cells were removed based on predefined filtering criteria, and subsequent steps were taken to correct for technical variations in the dataset according to best practices^59^. Specifically, total counts, the percentage of mitochondrial counts, and the percentage of External RNA Controls Consortium (ERCC) spike-in counts were regressed out to normalise the data and mitigate technical confounding effects. The preprocessing steps and scripts used are available at [dataset preprocessing and removal of low-quality cells](https://www.yizhouyu.com/opc2sc/scripts/0_1_dataset_formatting_invivo_counts.html) and [regressing out unwanted variables](https://www.yizhouyu.com/opc2sc/scripts/1_1_dataset_formatting_regressout.html). Following preprocessing, cell states, also referred to as cell types, were annotated. This annotation was performed using marker-based identification to ensure accurate classification of distinct cell populations. The full annotation workflow can be accessed via our [cell state annotation](https://www.yizhouyu.com/opc2sc/scripts/1_2_invivo_celltype_annodation_regressoutdataset.html) script.

To investigate the dynamics of OPCs in both injured and control conditions, cell trajectory analysis was conducted using Partition-based Graph Abstraction (PAGA)^60^. This approach allowed for the reconstruction of differentiation trajectories, capturing the progression of OPC states across conditions. The full analysis pipeline for PAGA trajectory inference is available at [PAGA trajectory analysis](https://www.yizhouyu.com/opc2sc/scripts/2_1_invivo_paga_analysis_wholedata.html). In addition to trajectory inference, cell fate mapping was performed using CellRank, which employs probabilistic modeling to infer the future states of cells based on transcriptomic similarities. The methodology for cell fate inference using CellRank is detailed at [cell fate mapping via CellRank](https://www.yizhouyu.com/opc2sc/scripts/2_2_invivo_paga_cellrank_wholedata.html). Transcription factor enrichment analysis was performed using ChEA3^61^ and pathway enrichment using STRING^62^.

For both BMP-treated OPCs and those subjected to Sox10 overexpression or Olig2 shRNA knockdown, preprocessing steps included quality control, filtering of low-quality cells, and normalization of gene expression counts using similar pipelines ([BMP-treated QC](https://www.yizhouyu.com/opc2sc/0_3_invitro_BMP4_QC.html), [TF-perturbed QC](https://www.yizhouyu.com/opc2sc/0_4_invitro_TF_QC.html)). To assess transcriptional dynamics, RNA velocity analysis was performed using scVelo to infer the future states of cells based on spliced and unspliced mRNA kinetics ([BMP-treated RNA velocity](https://www.yizhouyu.com/opc2sc/3_3_1_invitro_cellrank_BMP4.html), [TF-perturbed RNA velocity](https://www.yizhouyu.com/opc2sc/3_3_2_invitro_cellrank_TF.html)). Cell fate mapping was then conducted with CellRank to predict differentiation trajectories influenced by BMP signalling or transcription factor perturbations ([BMP-treated CellRank](https://www.yizhouyu.com/opc2sc/3_3_1_invitro_cellrank_BMP4.html), [TF-perturbed CellRank](https://www.yizhouyu.com/opc2sc/3_3_2_invitro_cellrank_TF.html)).

For the analysis of human multiple sclerosis snRNA data, count matrices from Lerma-Martin and colleagues^18^ were downloaded. Nuclei labelled as OPCs by the original authors were extracted, preprocessed, and re-integrated using scVI^63^ the donor, number of genes by counts and percentage of mitochondrial counts as continuous variables. For the identification of oSCs in human OPCs, we first trained a model using CellTypist^64,65^ the rat demyelination single cell data, and used label transfer to predict the putative oSC in samples from human MS patients. Pathway and transcription factor analyses were performed as described above, and cell-cell communication analysis was done using CellPhoneDB^19^.

### Statistical analysis

Statistical analyses were performed using R and GraphPad Prism. Prior to data analysis, Shapiro-Wilk tests were performed to confirm normality of the data. The data are presented as the mean values, and the error bars indicate ± standard deviation (s.d.). The number of biological replicates per experimental variable (n) is indicated in the respective figure legend. Blinding was not performed. Significance is indicated as *p<0.05, **p<0.01, ***p<0.001, ****p<0.0001 and not significant (n.s.) for p>0.05.

---

**Extended Data Fig. 1:**
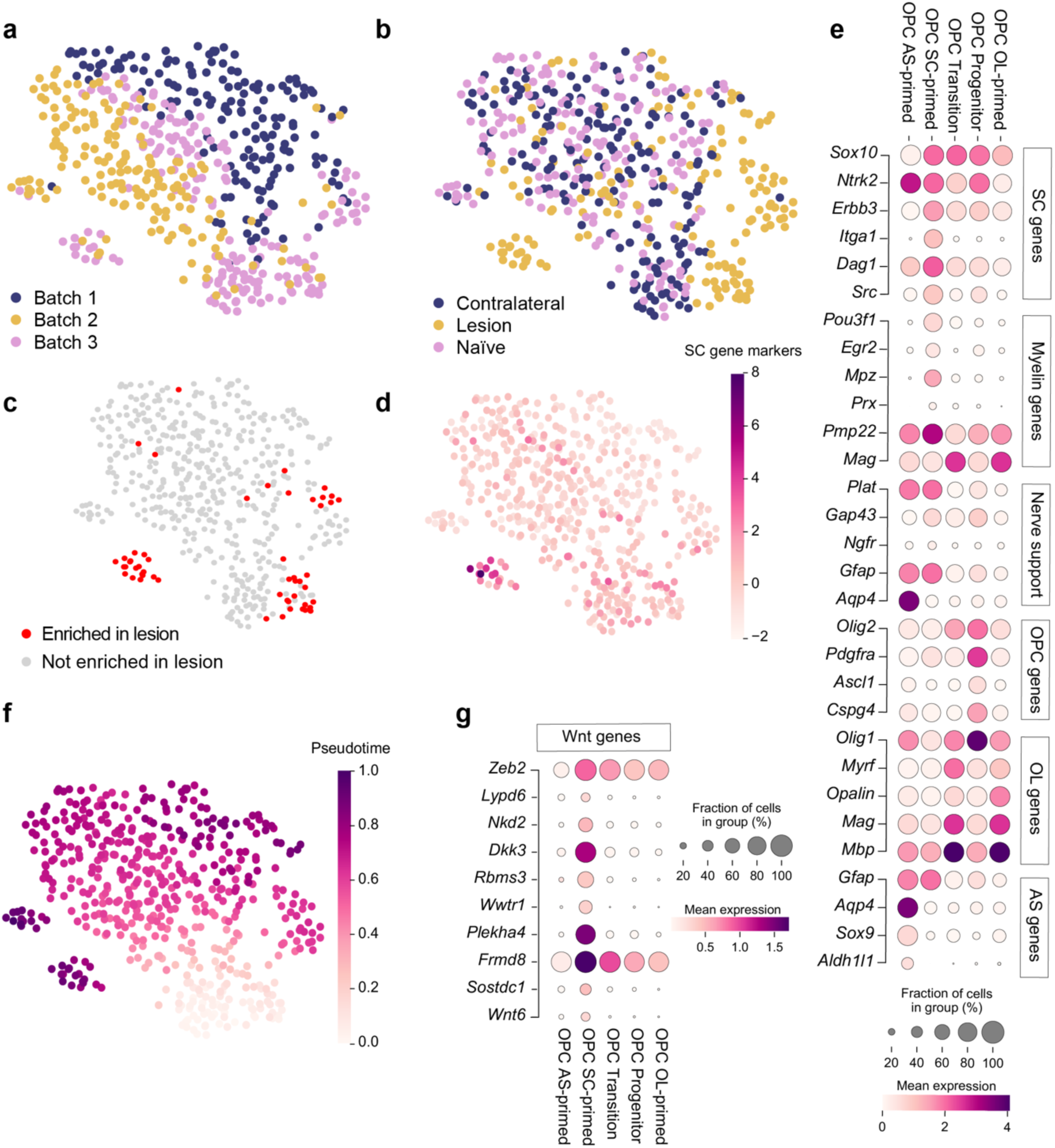
Transcriptomic analyses confirm data robustness and reveal that SC-primed OPCs are unique to the lesion. **a, b**, UMAP visualisation stratified by batch (**a**) and by condition (**b)**. **c,** Differential abundance analysis based on K-nearest neighbours using Milo. Cells from clusters that are significantly increased (spatial FDR<0.05) in the injury condition are labelled in red. No cell clusters were significantly decreased in this comparison. **d,** Expression of a SC gene score, calculated using a list of SC-related genes (*Mpz, Prx, Egr2, Sox10, Pou3f1*) subtracted by the average expression of a random set of reference genes. **e,** Dot plot showing extended marker gene expression across different OPC sub-populations. The analysis includes genes associated with SC identity, myelination, nerve support, and CNS glial cell lineages (OPCs, oligodendrocytes, and astrocytes). **f,** Trajectory analysis to predict the differentiation pseudotime using graph abstraction. **g,** Expression of a set of fate driver genes identified in Fig. 1e using CellRank that are related to the Wnt pathway.

**Extended Data Fig. 2:**
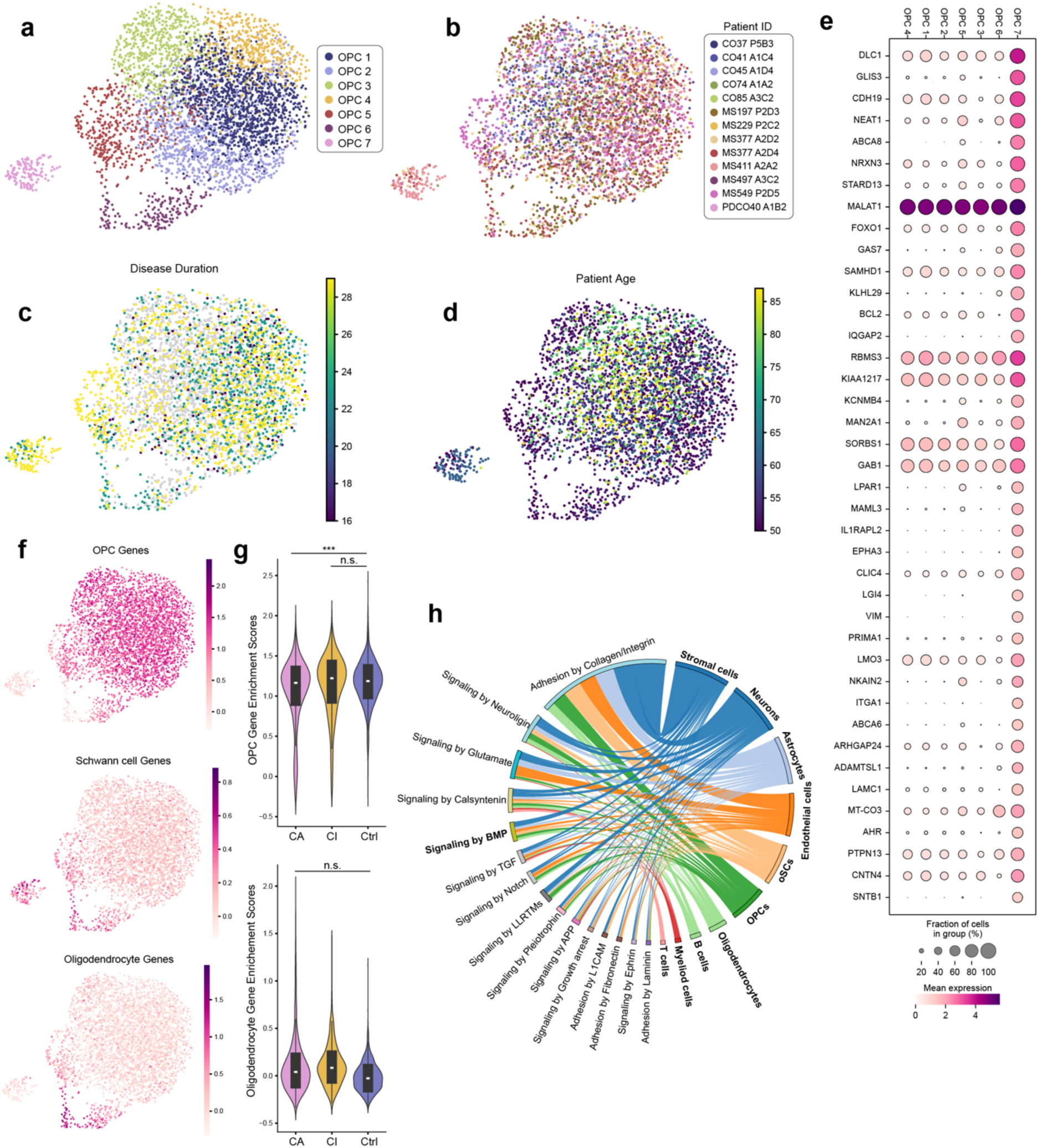
Extended characterisation of SC-primed OPCs in human MS. **a-d**, UMAP projection of OPCs from the Lerma-Martin et al., 2024 dataset with cells coloured by (**a**) Leiden clusters, (**b**) patient ID, (**c**) disease duration (in years, grey dots correspond to nuclei where that data was not available) and (**d**) patient age. **e,** Top differentially expressed genes from the predicted SC-primed OPCs (Cluster 7) across the seven OPC subclusters. Significance was determined using the Wilcoxon rank sum test with FDR adjustmexnt. **f,** UMAP projections of OPCs, with cells coloured by their enrichment scores for lineage gene signatures of OPCs (Top), SCs (Middle) and OLs (Bottom). The list of marker genes used to calculate the enrichment scores is described in Extended Data Table 1. **g,** Violin plots showing the scores of OPC and oligodendrocyte gene signatures, stratified by disease condition: chronic inactive (CI), chronic active (CA), and control. **h,** Ligand-receptor interactions between oSCs and other cell types in the brain. The class of signalling pathways is on the left, and cell types on the right are highlighted in bold.

**Extended Data Fig. 3:**
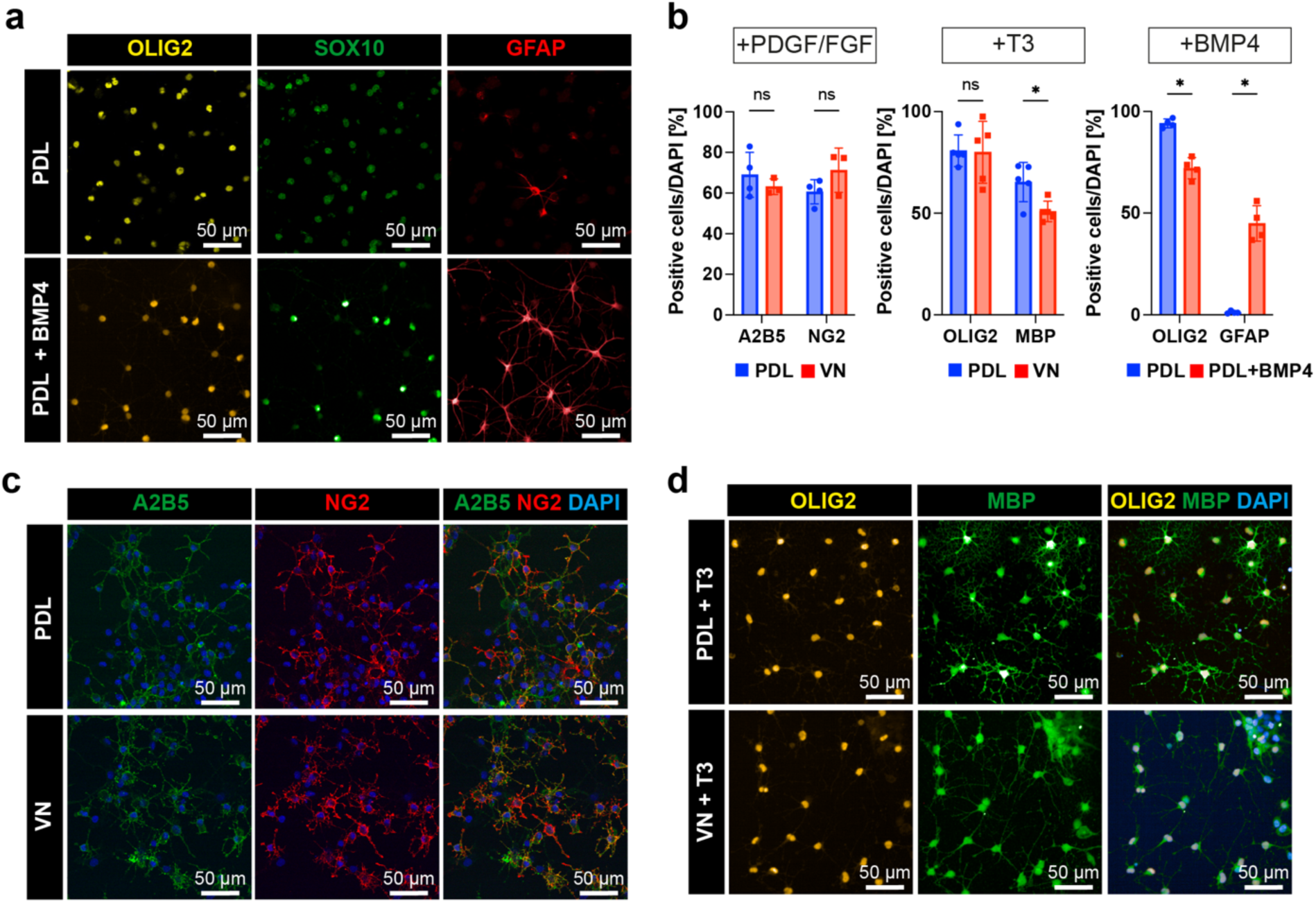
Vitronectin or BMP4 alone cannot induce OPC-to-SC differentiation. **a**, OLIG2, SOX10, and GFAP immunostaining of OPCs cultured on PDL with or without BMP4. **b,** Quantification of cells immunostained for A2B5, NG2, OLIG2, MBP, or GFAP under the specified conditions. Values are mean ± SD, (*p< 0.05, **p< 0.01 by unpaired t-test; ns, not significant). Data from at least three independent experiments. **c,** A2B5 and NG2 immunostaining of OPCs cultured on PDL or vitronectin under proliferation conditions. **d,** OLIG2 and MBP immunostaining of OPCs cultured on PDL or vitronectin under differentiation conditions.

**Extended Data Fig. 4:**
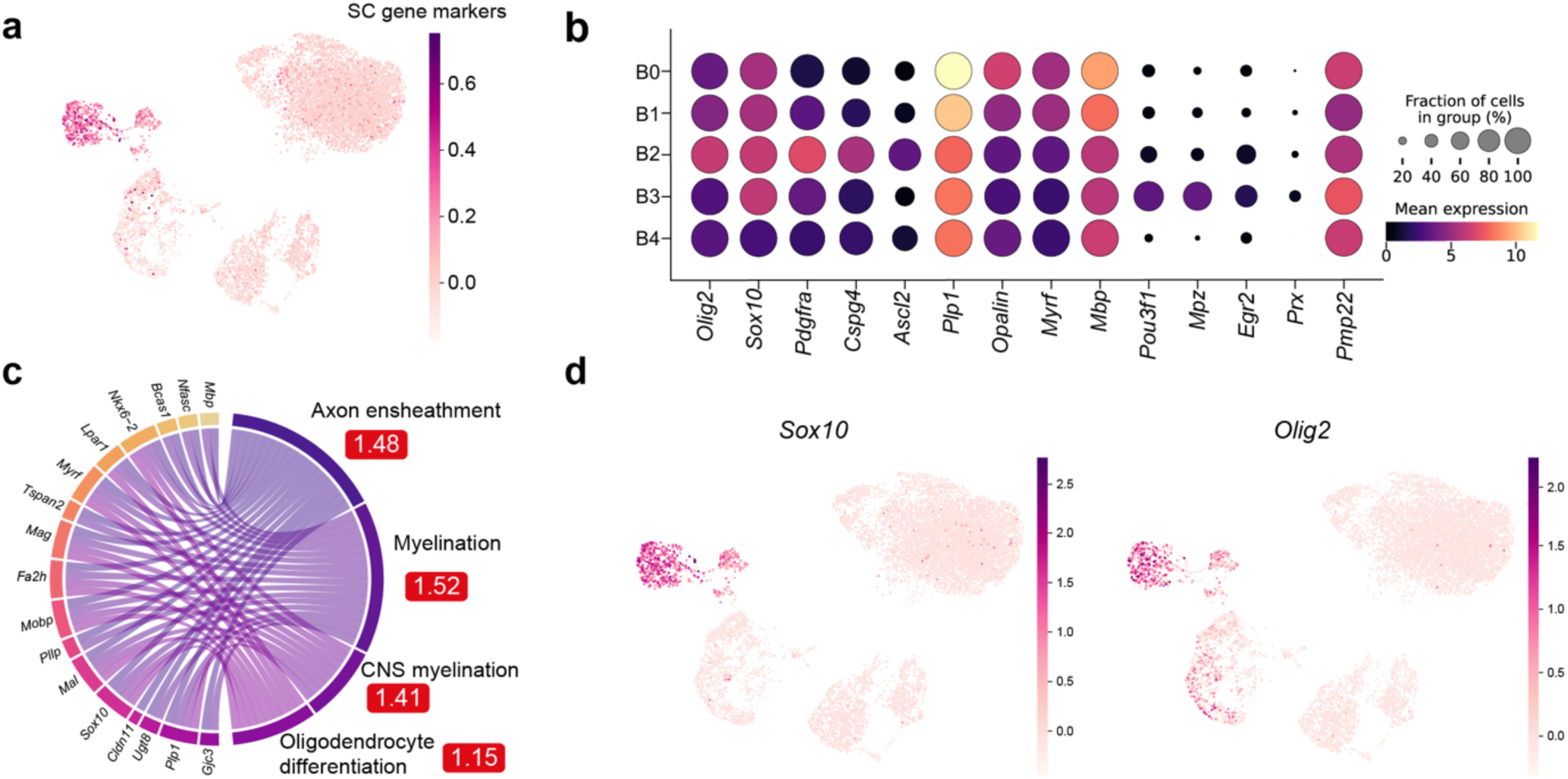
Extended characterisation of Vn/BMP4-induced oSCs. **a**, Expression of SC gene score composed of SC genes including (*Mpz*, *Prx*, *Egr2*, *Sox10* and *Pou3f1*) in oSCs treated with Vn/BMP4. **b,** Expression dotplot of key lineage and functional marker genes across the five Vn/BMP4 treated OPC subclusters. **c,** Pathway enrichment analysis of the top 200 fate driver genes identified using CellRank2 and their corresponding pathways. The enrichment scores are shown in red boxes. Only a subset of pathways are shown; an extended list is available here: https://izu0421.github.io/opc2sc. **d,** Expression of *Sox10* and *Olig2* in Vn/BMP4-induced oSCs.

**Extended Data Fig. 5:**
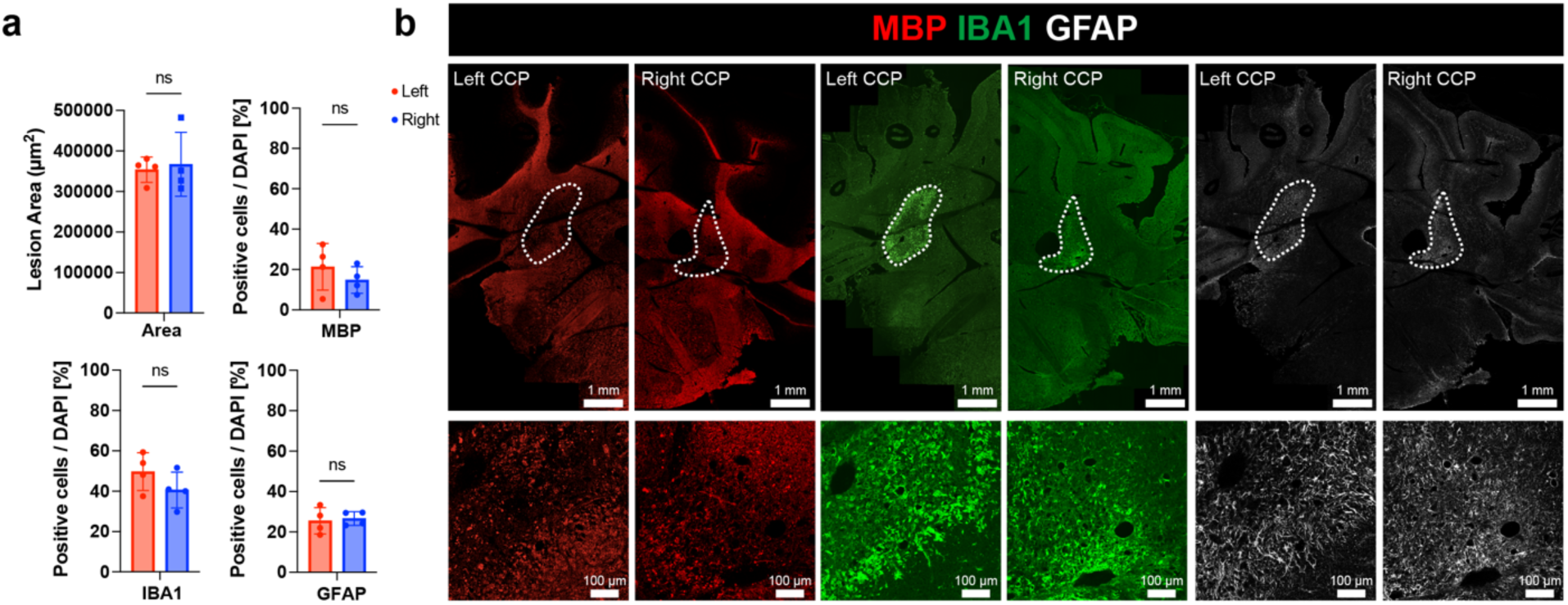
Transplanted BMP4/Vn-induced SCs remyelinate CNS axons, without perturbing host lesion dynamics. **a**, Quantification of the lesion area, and the number of MBP, IBA1, and GFAP-positive cells within the lesion. **b,** MBP, IBA1, and GFAP immunostaining to visualise the bilateral lesion and unilateral transplant area at 28 dpl

**Extended Data Fig. 6:**
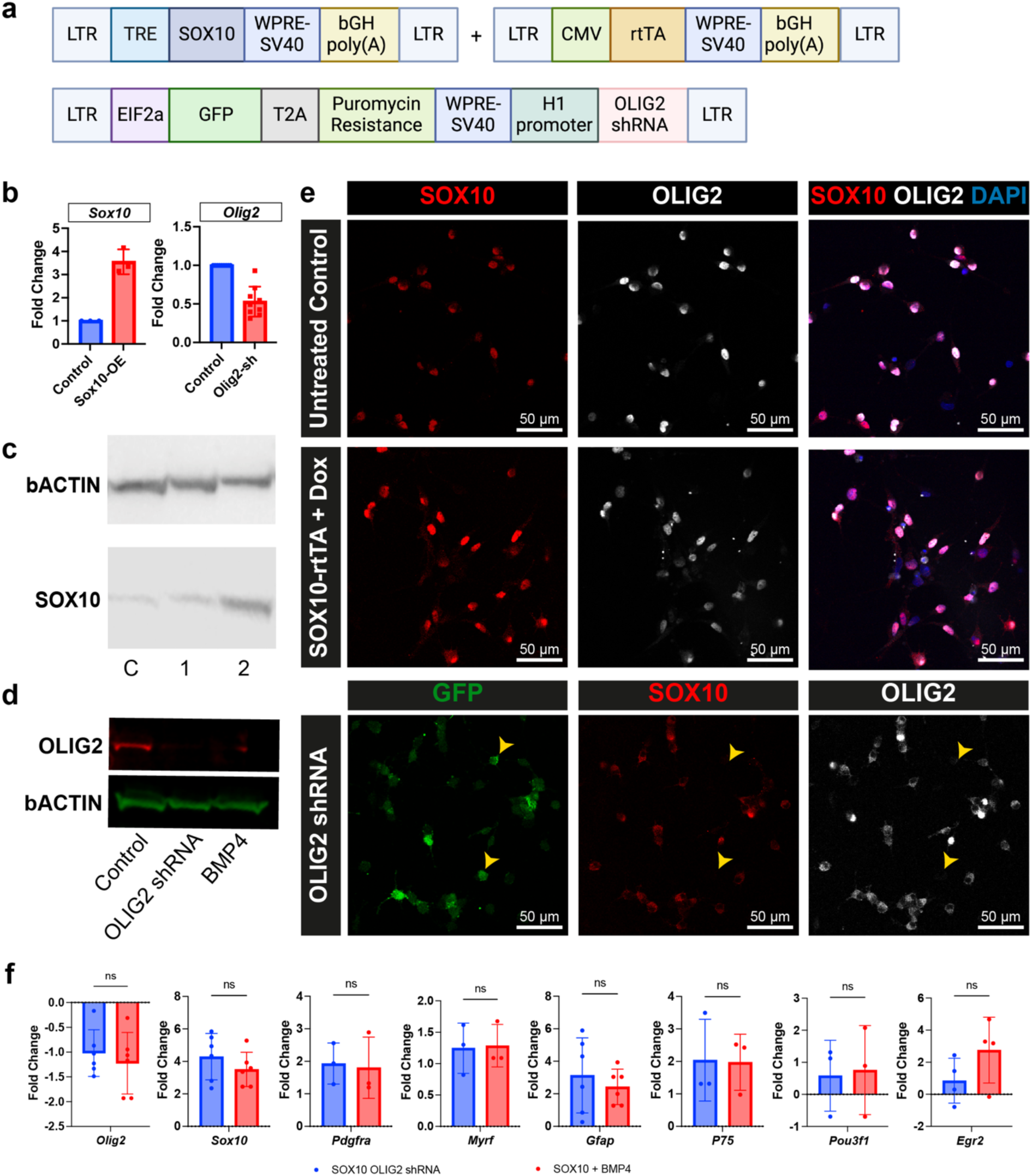
Generation and validation of constructs to overexpress Sox10 and to knockout Olig2. **a**, Schematic linear map of the LV-Sox10, LV-rtTA and LV-OLIG2shRNA plasmid constructs. **b,** RT-qPCR analysis of *Olig2* and *Sox10* mRNA expression in cells infected with LV-Sox10, or LV-Olig2 shRNA, compared to uninfected controls. Values are presented as mean ± SD. (*p< 0.05, **p< 0.01, ***p< 0.001, ****p< 0.0001 by unpaired t-test; ns, not significant). **c,** Western blot showing SOX10 protein levels in OPC lysates. Lanes represent: (C) untreated controls, (1) cells infected with LV-TRE-Sox10 only, and (2) cells co-infected with LV-TRE-Sox10 and LV-CMV-rtTA. **d,** Western blot showing OLIG2 protein levels in OPCs treated with LV-Olig2shRNA compared to untreated and BMP4-treated controls. **e,** SOX10 immunostaining in OPCs infected with LV-TRE-Sox10 and LV-CMV-rtTA compared to untreated controls. The figure also includes images of GFP^+^ cells, which were infected with LV-OLIG2 shRNA, showing the fluorescence intensity of SOX10 and OLIG2. Data are representative of three independent experiments. **f,** RT-qPCR analysis of oligodendroglial genes (*Olig2*, *Sox10*, *Pdgfra*, *Myrf*) and SC lineage genes (*Gfap*, *p75*, *Oct*, *Krox20*) in OPCs treated with LV-Sox10 and LV-OLIG2shRNA versus LV-Sox10 and BMP4. Values are mean ± SD, (*p< 0.05, **p< 0.01, ***p< 0.001, ****p< 0.0001 by unpaired t-test; ns, not significant).

**Extended Data Fig. 7:**
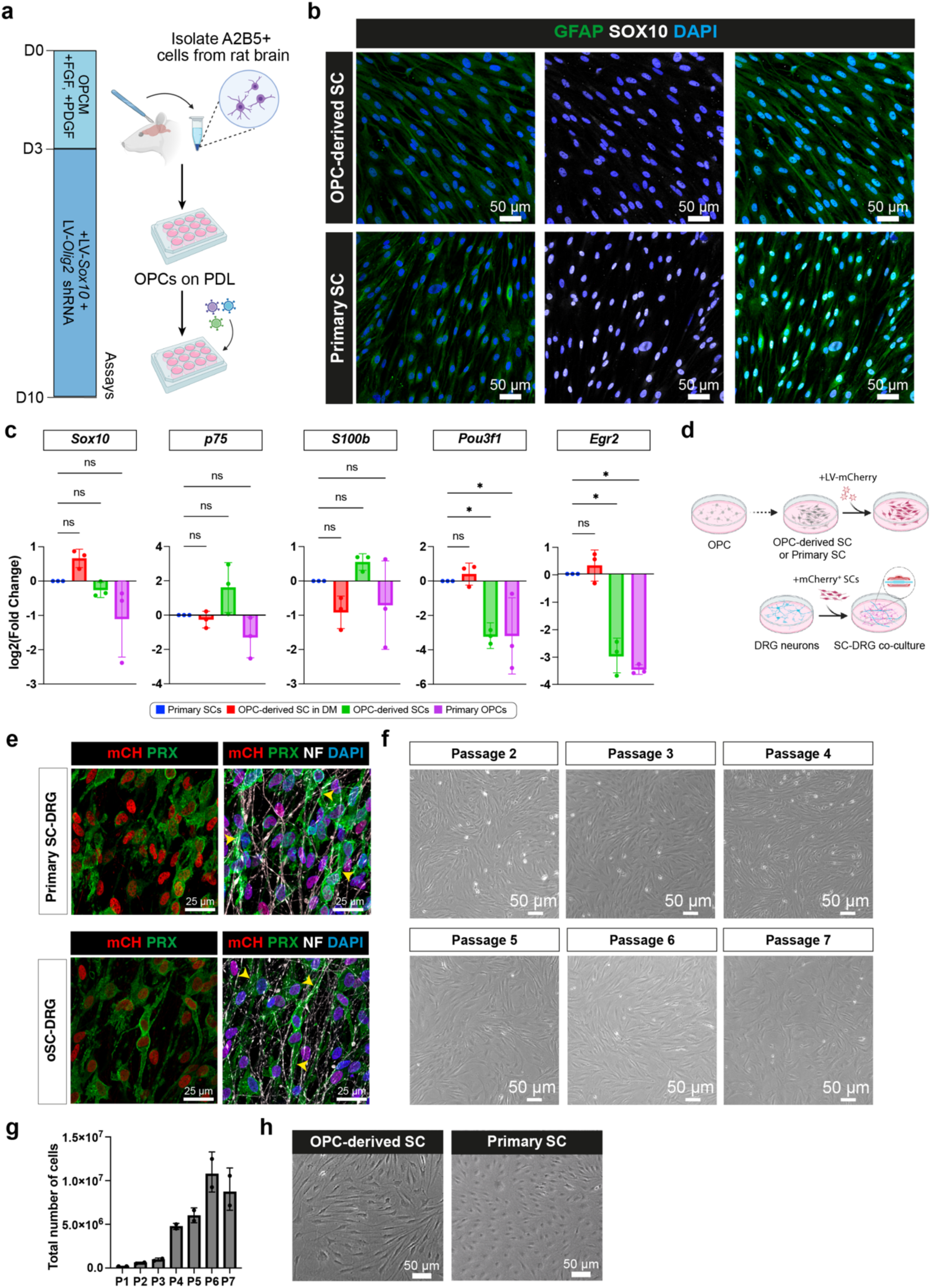
Suppressing Olig2 and stabilising Sox10 activity is sufficient to induce OPCs to convert into Schwann cells *in vitro*. **a**, Induction of OPC-to-SC differentiation. OPCs were isolated from an adult rat brain, then maintained in OPC proliferation medium (+FGF, +PDGF) to allow cells to recover for three days. Lentiviruses encoding *Sox10*, *rTTA*, and *Olig2* shRNA were then added to infect OPCs, which were subsequently cultured until day 10 (D10). **b,** Immunostaining for GFAP and SOX10 in oSC and primary SC cultures. **c,** RT-qPCR analysis of myelinating SC markers (*Sox10*, *Pou3f1*, and *Egr2*), and immature/non-myelinating SC markers (*p75* and *S100b*) in primary OPCs, oSCs, and primary SCs. The bar graph shows mean gene expression ± SD. Gene fold change is normalised to primary SCs. Data are presented as mean ± SD. Statistical significance was determined by Kruskal-Wallis test followed by Dunn’s multiple comparisons test; *p< 0.05, **p< 0.01, ***p< 0.001, ****p< 0.0001, ns, not significant). Each group included at least three biological replicates. **d,** Schematic showing SC-DRG co-culture experiment. SCs were pre-labeled with LV-mCherry before being seeded onto DRGs to exclude the possibility of SC contaminants in DRG preparations. **e,** Immunostaining of PRX and mCherry in myelinating co-cultures of DRGs with either mCherry-labelled oSC or primary SCs. Data are representative of at least three independent experiments. **f,** Representative bright-field images of oSCs at different passages (P2 to P7). **g,** Total number of SOX10/OLIG2-induced oSCs from passage 2 (P2) to passage 7 (P7). Values are mean ± SD. **h,** Bright-field images of oSCs and primary SCs.

**Extended Data Fig. 8.**
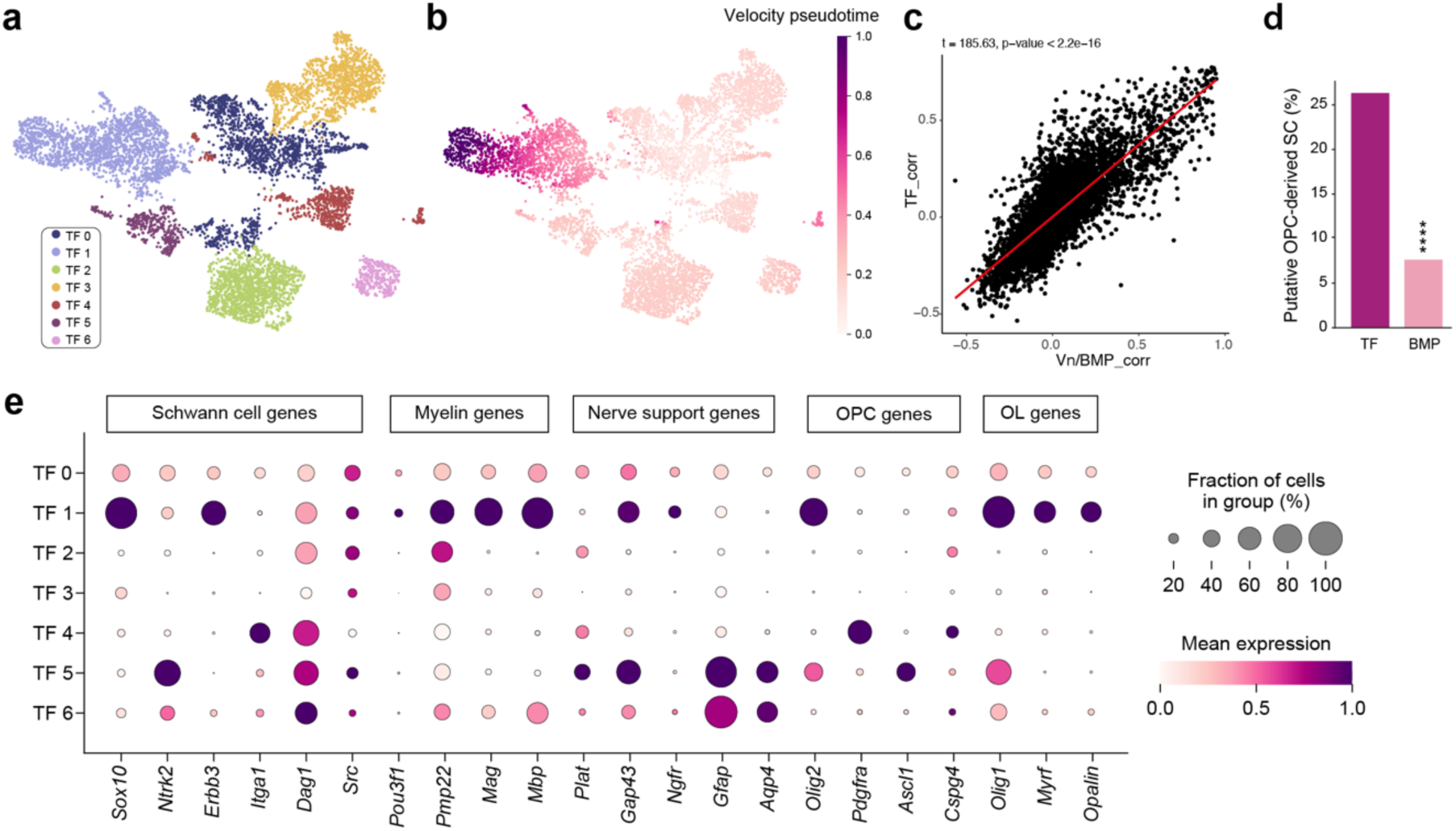
Extended characterization of OPC-derived Schwann cells. **a**, Total number of SOX10/OLIG2-induced oSCs from passage 2 (P2) to passage 7 (P7). Values are mean ± SD. UMAP representation of scRNA-seq data from the SOX10/OLIG2 -induced oSCs reveals seven cellular clusters. **b,** RNA velocity analysis of SOX10/OLIG2-induced oSCs. **c,** Correlation between SC cell fate driver genes in the SOX10/OLIG2-induced (y axis) and BMP/Vn-treated (x axis) datasets (Pearson’s product-moment correlation, t = 185.63, p-value < 2.2e-16). Each dot represents a gene. **d,** Bar graph showing the percentage of oSCs from BMP4/Vn and SOX10/OLIG2 -treated OPCs out of the total number that passed quality control and were sequenced. **e,** Expression of SC lineage-associated transcription factors and functional markers across the seven cell clusters.

**Extended Data Fig. 9:**
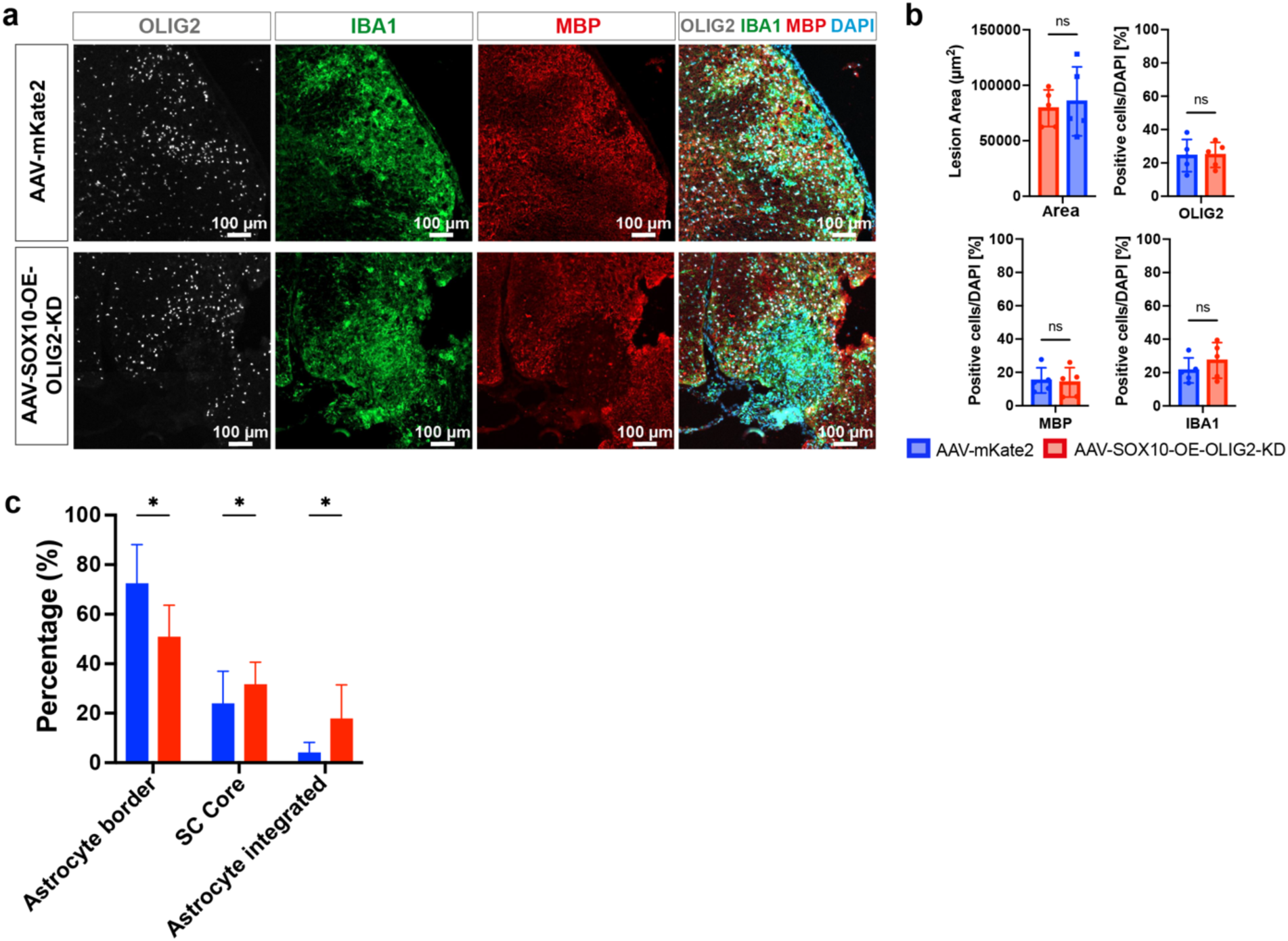
AAV induction does not alter lesion dynamics following demyelination. **a**, Immunostaining of OLIG2, IBA1 and MBP in the lesion area. **b,** Quantification of the data presented in (**a**) The bar graphs show the lesion size and the number of OLIG2^+^, MBP^+^, and IBA1^+^ cells in animals receiving AAV-SOX10/OLIG2 or AAV-mKate2-treated animals. Values are expressed as mean ± SD. Statistical significance was determined by one-way ANOVA with Tukey’s multiple comparisons test. **c,** Quantification of the spatial relationship between PRX^+^ SCs and GFAP^+^ astrocytes in the lesion site.

## Extended Data Tables

**Extended Data Table 1:**
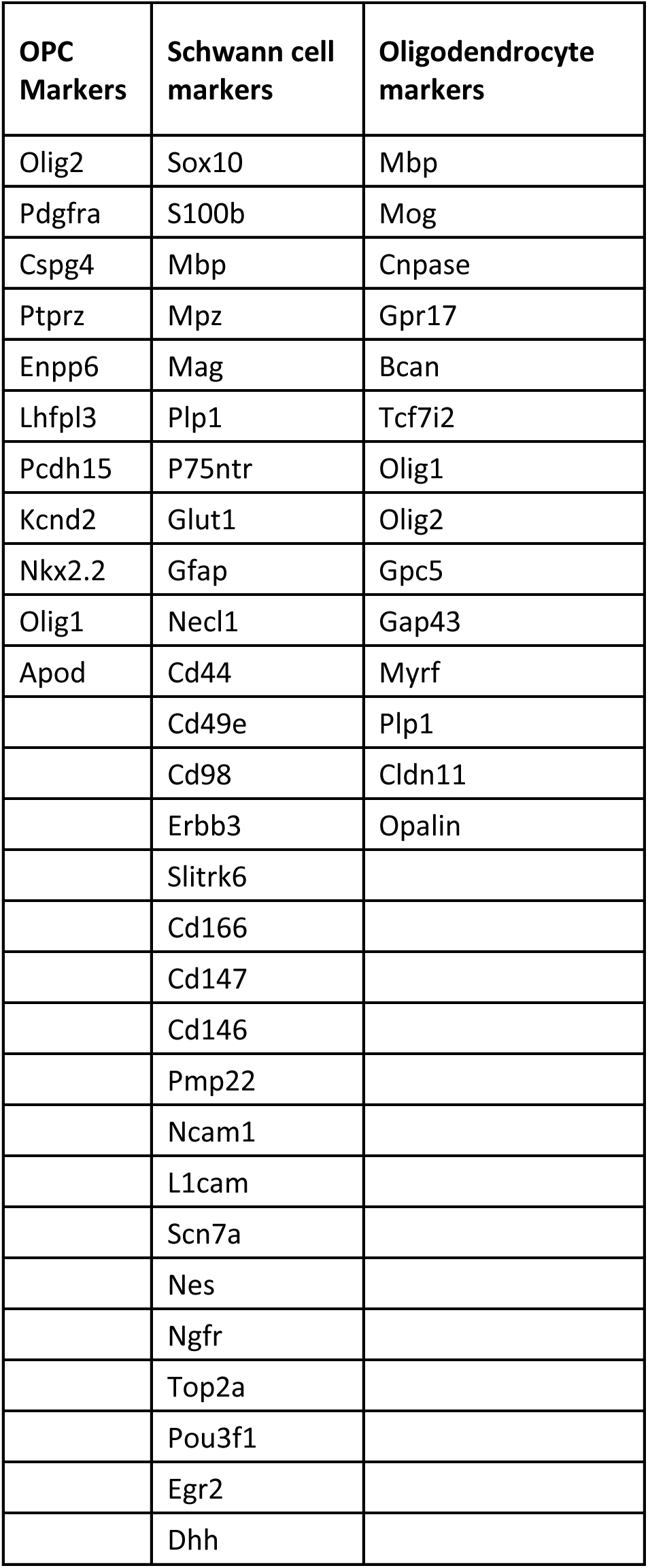
List of genes used to calculate OPC, SC and Oligodendrocyte enrichment scores.

**Extended Data Table 2:** List of antibodies used in this study.

| **Antibody Name** | **Catalogue Number** | **Company** | **Host** | **Dilution** |
| --- | --- | --- | --- | --- |
| A2B5 | MAB312 | Millipore | Mouse | 1:1000 |
| bACTIN | AB6276 | Abcam | Mouse | 1:500 |
| B-III-Tubulin | MMS-435P | Biolegend | Mouse | 1:1000 |
| CC1 | MABC200 | Sigma | Mouse | 1:500 |
| GFAP | AB46744 | Abcam | Chicken | 1:2000 |
| HA | 26183 | Invitrogen | Mouse | 1:500 |
| IBA1 | AF5076 | Abcam | Goat | 1:500 |
| MAP2 | AB5392 | Abcam | Mouse | 1:1000 |
| MBP | MCA4095 | Serotec | Rat | 1:200 |
| mCherry | AB125096 | Abcam | Mouse | 1:500 |
| NG2 | AB5320 | Sigma-Aldrich | Rabbit | 1:500 |
| OLIG2 | AB9610 | Abcam | Rabbit | 1:500 |
| OLIG2 | MABN50 | Sigma-Aldrich | Mouse | 1:500 |
| MPZ | AB134439 | Abcam | Chicken | 1:500 |
| PRX | HPA001868 | Cambridge Bioscience | Rabbit | 1:500 |
| SM132 | 801701 | Biolegend | Mouse | 1:1000 |
| SOX10 | AF2864 | R&D | Goat | 1:200 |
| SOX10 | AB155279 | R&D | Rabbit | 1:500 |

**Extended Data Table 3:**
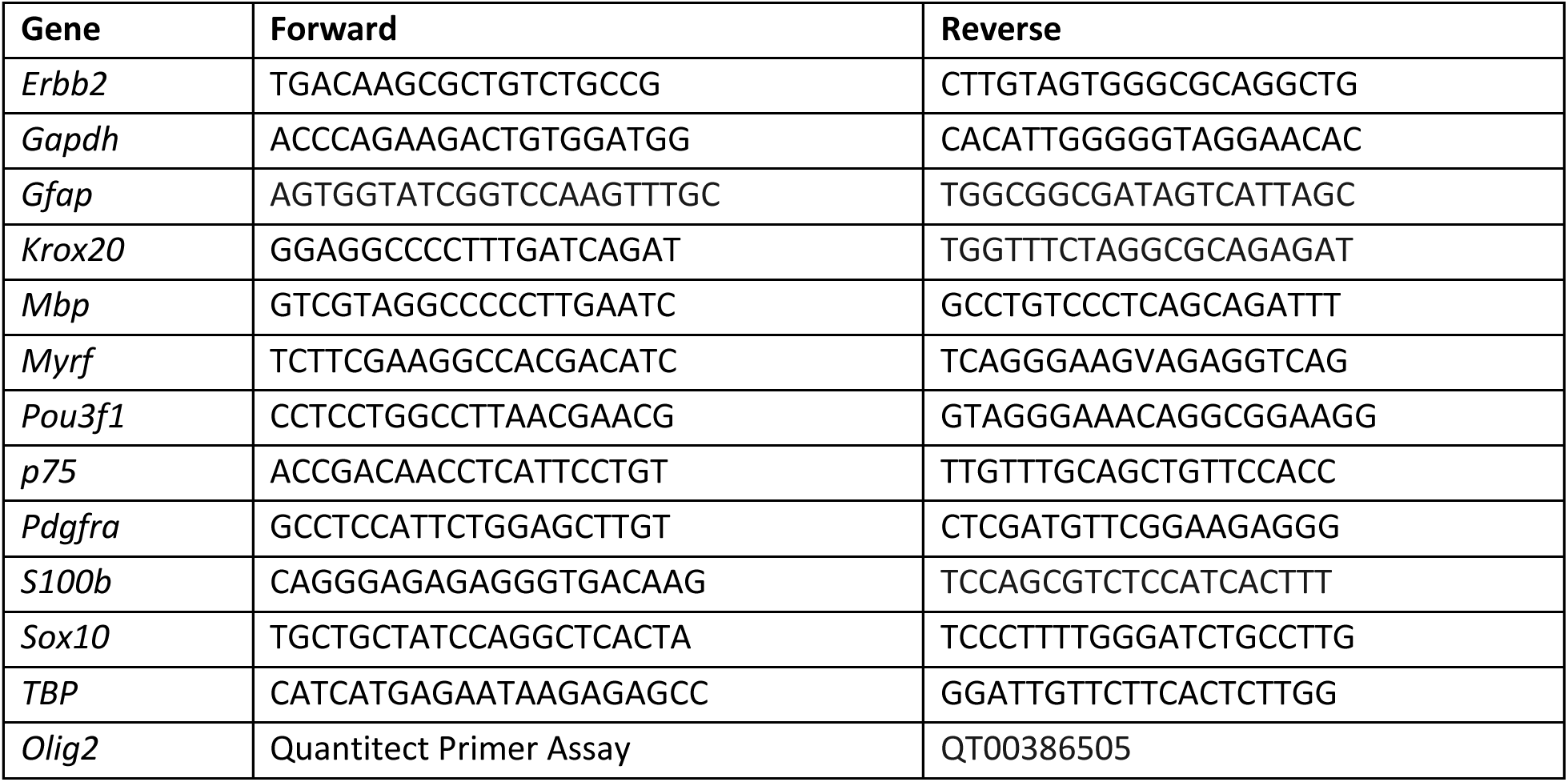
List and sequence of qPCR primers used in this study.

